# Unraveling Glioblastoma Heterogeneity: Introducing SP2G Method for Identifying Invasive Sub-Populations

**DOI:** 10.1101/2024.01.10.574982

**Authors:** Michele Crestani, Nikolaos Kakogiannos, Fabio Iannelli, Tania Dini, Claudio Maderna, Monica Giannotta, Giuliana Pelicci, Paolo Maiuri, Pascale Monzo, Nils C. Gauthier

## Abstract

Glioblastomas exhibit remarkable heterogeneity at various levels, including motility modes and mechanoproperties that contribute to tumor resistance and recurrence. In a recent study using gridded micropatterns mimicking the brain vasculature, we linked glioblastoma cell motility modes, mechanical properties, formin content, and substrate chemistry. We now introduce SP2G (SPheroid SPreading on Grids), an analytic platform designed to identify the migratory modes of patient-derived glioblastoma cells and rapidly pinpoint the most invasive sub-populations. Tumorspheres are imaged as they spread on gridded micropatterns and analyzed by our semi-automated, open-source, Fiji macro suite that characterizes migration modes accurately. With SP2G, we could reveal intra-patient motility heterogeneity with molecular correlations to specific integrins and EMT markers. Thus, our system presents a versatile and potentially pan-cancer workflow to detect diverse invasive tumor sub-populations in patient-derived specimens and offers a valuable tool for therapeutic evaluations at the individual patient level.

**Teaser:** Cracking the inter and intra-patient diversity in Glioblastoma migration profiles

## Introduction

Glioblastoma (GBM) tumors are highly aggressive and debilitating primary brain tumors that lead to the destruction of the brain and the rapid death of the patients (8-14 months median survival) (*1, 2*). To this day, GBMs remain incurable, even with aggressive treatments that include brain surgery, radio-and chemo-therapies (*1, 3, 4*).

The massive invasiveness and the inter- and intra-patient heterogeneity of GBM cells, represent two major clinical challenges in GBM treatment. High heterogeneity precludes distinguishing the most aggressive cells from the rest of the tumor and thus, defining precise and valuable therapeutic targets (*5–12*). Invasiveness allows cells to escape treatment and to generate secondary tumors responsible for the systematic recurrence observed in GBM patients (*13, 14*). Previous treatments targeting classical migratory pathways have failed to improve GBM outcome (*13, 15, 16*). This is probably because GBM cells use their own specific motility modes to invade the brain (*13, 14, 17*). In contrast to metastatic cells, glioma cells do not rely on the bloodstream for spreading. Instead, they engage in active migration along specific pathways, such as the abluminal surface of brain blood vessels and the white matter tracts, commonly referred to the ‘structures of Scherer’, and invade, albeit less efficiently, by navigating into the brain parenchyma (*18–32*). These diverse motility modes need to be inventoried and understood in order to find specific molecular targets to stop GBM cells from spreading into the brain.

Currently, the most effective biological tool for studying the cell biology of GBM is the utilization of cell lines derived from patient samples. These cells are able to grow and form a GBM tumor when injected into mouse brains (and hence are called human glioma propagating cells, hGPCs) (*33–35*). hGPCs are usually grown as spheroids (tumorspheres) in serum-deprived medium, allowing keeping their original properties including cell stemness and heterogeneity (*33–35*). Spheroids constitute a great tool to dissect the motility modes of cell escaping the tumor core. They have been used in many motility studies including those aiming at understanding the influence of biomaterials, microenvironmental cues, drugs, mostly in 3D systems that do not recapitulate the brain linear tracks exploited by GBM cells as invasive highways (*36–44*). Spheroids have also been used in brain slices and xenografts that fully recapitulate the brain linear tracks and allow the interconnection of invading GBM in communicating networks, but that require laborious protocols and are inappropriate for robust analytical tool development needed to accelerate discoveries and diagnosis (*38, 45–48*). Altogether, these impediments preclude a holistic dissection of GBM migration and motility modes and fail to provide a robust and broadly applicable platform to unveil how the cancer heterogeneity found in patients impacts on cell invasiveness through motility along brain topographical cues.

Bioengineered systems have been set up to recapitulate the brain linear tracks in controlled environments such as micropatterns, channels, nanofibers and grooves (*49–52*) and that promote glioma motility, especially if laminin is used (*53–57*). Nanofibers and linear grooves recapitulate well the 3D environments but are difficult to image at high resolution (*53, 54, 57, 58*). Engineered vessels and microvascular networks potentiate GBM migration but are inadequate for broad systematic investigation due to the co-culture setups that add complexity and compromise reproducibility (*49–51*). In previous studies, we demonstrated that linear micropatterns were excellent proxy to mimic the brain blood vessel tracks (*50, 56, 59*). These micropatterns allow high-resolution imaging and analysis, while recapitulating the stroke motion observed *in vivo*. They also present the advantages of simple in vitro systems, such as experimental reproducibility and relatively short experimental duration (*50, 56, 59, 60*).

Recently, we further improved the linear system with gridded micropatterns that recapitulate better the brain vasculature complexity and could reveal differences in motility that were not identifiable using lines only (*55, 61*). Using this system, two different motile behaviors were defined: hurdling and gliding. The hurdlers could ‘jump’ over the passivated area of the substrate and cut grid corners, while gliders were diligently following the tracks. We also found that hurdlers and gliders were mechanically different and displayed diverse formin expression profiles (*55*).

In the present study, we further improved our system by combining the gridded micropatterns with spheroid spreading assays (SP2G for SPheroid SPreading on Grids) and tailored an ImageJ/Fiji toolbox able to translate the imaging data into numerical outputs reflecting the migration efficiency of the cells and their motility modes. This system allows a fast and semi-automatized analysis and can rapidly detect heterogeneous populations.

## Results

### SP2G: a bona fide assay to mimic brain blood vessel motility

To perform SP2G assays, fluorescent glioma spheroids were seeded on gridded micropatterns and imaged immediately, allowing recording the first steps of the cells reaching linear substrates mimicking the blood vessel network. The features of the grids (7 µm width and 75 µm gaps) were chosen to match the blood vessel features and density found in the brain (Fig. S1C) (*61*). Grids were coated with laminin since this matrix protein is enriched around brain blood vessels and promotes single glioma cell motility (*50, 55, 56, 62–65*). Moreover, in accordance with previous studies (*66, 67*), we found that glioma spheroids spread faster on laminin than on other substrates (Fig. S1A-B). To evaluate the efficiency of our system, we compared SP2G with several conventional assays. Rat C6 glioma cells, a well-established model that migrate efficiently on brain vasculature (*22, 24, 25, 28*), were cultured as spheroids, stained and seeded on mouse brain slices, in 3D gels (collagen and reconstituted basement membrane (rBM)), on laminin-coated dishes (2D-flat) and on our gridded-micropatterns (Fig. 1). Cells were imaged immediately after seeding (movie 1) and the spheroid spreading was quantified by measuring the areas of the spheroids at different time points. As observed in fig. 1A-E, G, spheroids spread faster on 2D flat and gridded micropatterns (∼8h for the complete dissolution of the spheres) than on 3D gels (>24h) and brain slices (>48h). Single cell migration was quantified by tracking manually the cells escaping the spheroids and calculating mean square displacement and mean velocity. Cells migrated faster in the setups providing linear guidance (brain slices and grids) compared to the other conditions (Fig. 1H, S1F-G). Moreover, on grids and brain slices, cells displayed elongated shapes and ‘stick-slip’ motility features (Fig 1I-J) (*22, 55, 56, 59, 68*), while in 3D gels cells protruded multiple finger-like structures, likely due to the tangled architecture of the microenvironment (Fig. 1I, arrowheads). On 2D flat, cells adopted a fan-like shape as previously described on homogeneous substrates (*56, 68*). Moreover, we found that spheroid spreading and single cell migration were independent of the spheroid original size, highlighting the flexibility of our assay since spheroid projected area can be highly variable ranging from 1,000 µm^2^ to 15,000 µm^2^ for the rat C6 (Fig. S1H-I). Overall, SP2G appeared as the closest decoy to recapitulate the conditions of a cell leaving the tumor core and encountering blood vessels. We compared SP2G with the other techniques for important parameters in GBM motility analysis, such as duration of the experiment (gain of time), presence of linear topographic cues, experimental reproducibility, optical accessibility, possibility to implement semi-automated analysis, 3D confinement and *in vivo* mimicry. SP2G appeared as an optimal technique in term of gain of time, optical accessibility and presence of linear topographic cues allowing stick-slip motility (Fig. 1F).

**Fig. 1.**
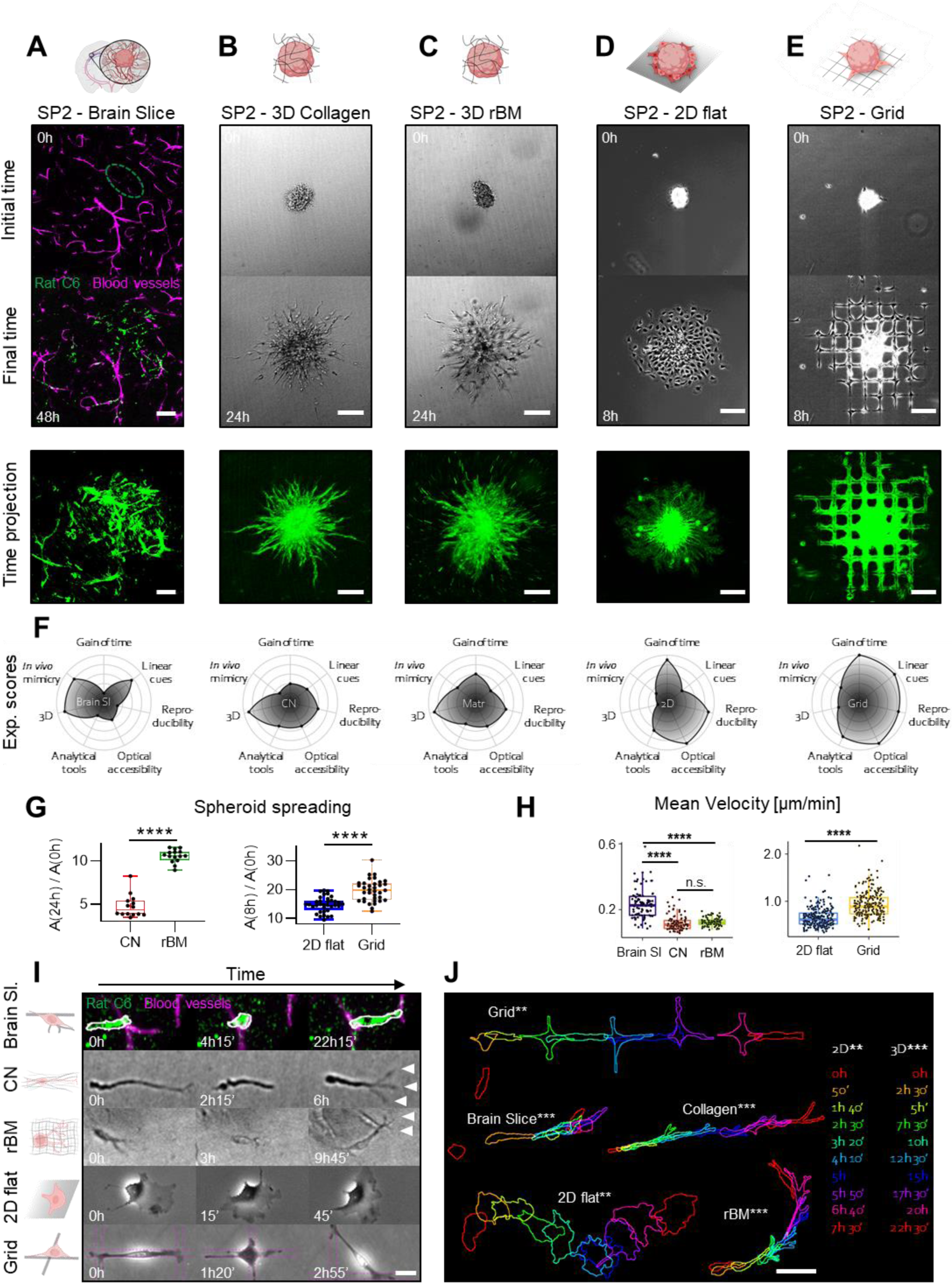
Spheroid spreading on grids (SP2G) mimics glioblastoma invasion on brain blood vessels. (**A-E**) Rat C6 glioma cells cultured as spheroids were stained (green, DiOC6 dye) and seeded in brain slices (**A**), in collagen gel (**B**), in reconstituted basement membrane (rBM) (**C**), on laminin-coated dishes (**D**), on gridded micropatterns (**E**) and imaged for 48h (**A**), 24h (**B, C**) or 8h (**D, E**). First and last images of the movies (upper panels) and time-projections (lower panels) are shown. The dashed oval in (**A**) corresponds to the initial area of the spheroid in the brain slice. (**F**) Radar plots summarizing experimental scores (1 to 5) of gain of time, presence of linear cues, experimental reproducibility, optical accessibility, possibility to develop analytical tools, three-dimensionality (3D) and *in vivo* mimicry for each setting. (**G**) Quantification of spheroid spreading in collagen gel, rBM, 2D flat and grid (n = 14, 15, 35, 35 spheroids respectively). (**H**) Mean velocities of single cells migrating in brain slices, collagen gel, rBM, 2D flat and grid (n = 80, 95, 90, 215, 215 tracks respectively, 5 to 7 tracks per spheroid, each dot is a cell). (**I**) Snapshots of single cells moving away from the spheroid in each setting, extracted from movie S1. (**J**) Panel summarizing cell shapes for C6 cell motility in each setting. Time is color-coded as indicated. Bars are 100 μm (**A-E**), 20 μm (**I**), and 50 μm (**J**).

### SP2G experimental setup and image analysis workflow

To quantitatively describe cell migration and motility modes, we designed a toolbox relying on 7 macros that deliver 3 outputs for cell migration (area, diffusivity, boundary speed) and 3 outputs for the motility modes (collective migration, directional persistence, hurdling) (Fig. 2A,B, see Supplementary Appendix for computational details and user manual). Since area measurements, A(t), neglect the temporal dimension, we integrated into SP2G diffusivity and boundary speed. Diffusivity, D(t), accounts for how quickly cells fill up the surrounding space. Boundary speed, V(t), indicates how quickly cells travel in a monodimensional space, providing a value equivalent to single cell velocity. Essentially, this toolbox allows translating migration modes and efficiency into unbiased numerical outputs, thus, defining a proper ‘motility signature’. For a better cell segmentation and a stable readout, grids and spheroids were stained with fluorescent dyes and spreading was imaged by fluorescence and phase contrast microscopy (Fig. 2A and Supplementary Video 2). We divided SP2G analysis workflow in 2 main steps: the first step processes the raw data semi-automatically and characterizes cell migration (outputs #1, 2, 3, Fig. 2C) and the second step characterizes motility modes (outputs #4, 5, 6, Fig. 2C). Cells and grid images are segmented in binary images that are multiplied to isolate grid nodes covered by the invasive cells. These binary images are combined to construct a polygon that connects the grid nodes travelled by the spheroid invasive front at each time point. An average migration area A(t) per cell line (>10 spheres per cell line) is represented by aligning the barycenter of all the polygons at time 0 and extrapolating all the mean xy coordinates at each time point (Fig. 2C, output #1). The corresponding numerical values are smoothed and differentiated to obtain diffusivity values D(t) (Fig. 2C, output 2). A(t) and D(t) values are then used to obtain the boundary speed v(t), from the formula v(t) = D(t) / (2√(A(t)) (output #3) (see Methods and Supplementary Appendix).

**Fig. 2.**
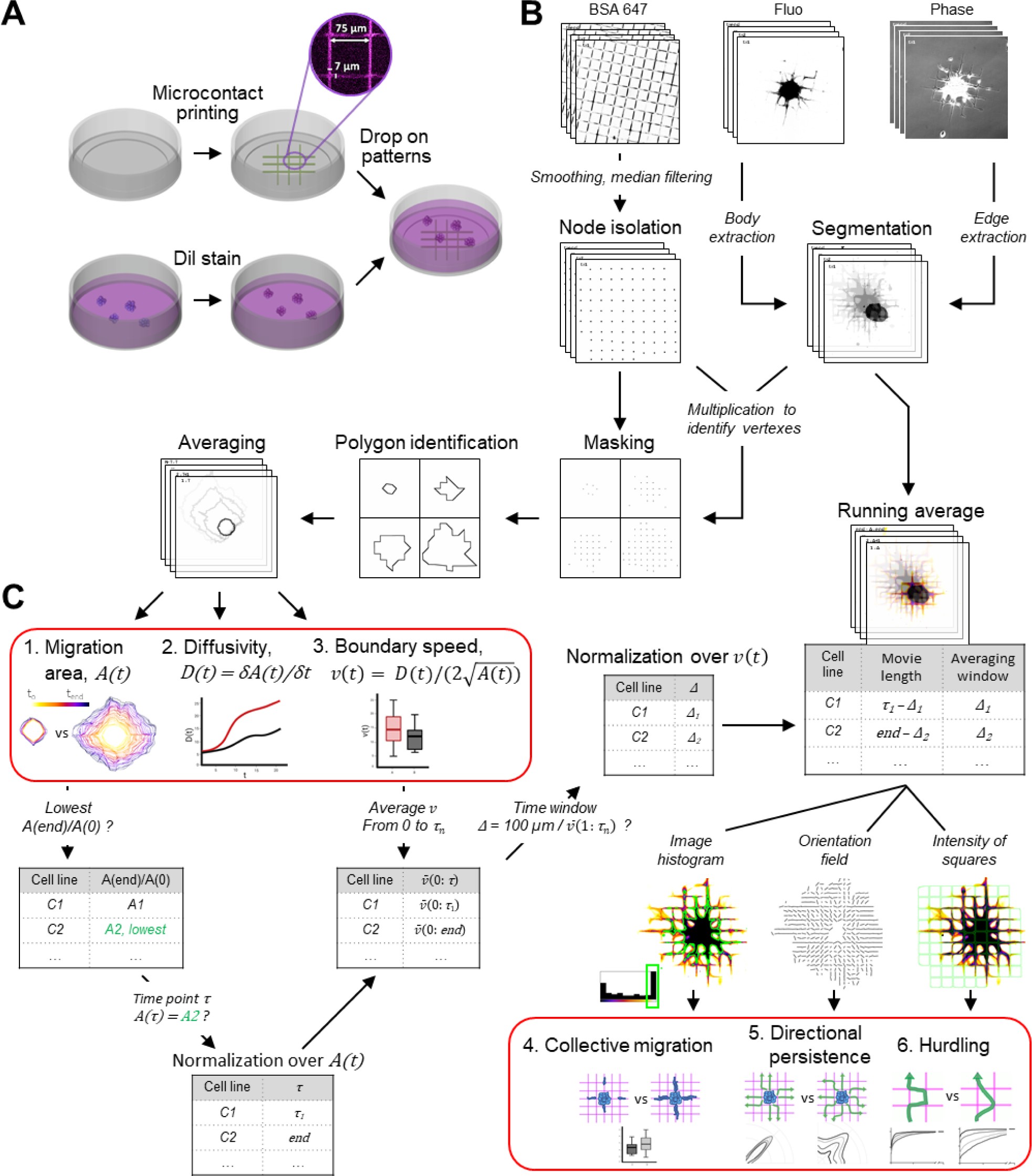
SP2G experimental setup and image analysis workflow. (**A**) Spheroids are stained (red, Dil), seeded on fluorescent gridded micro-patterns (coated with laminin mixed with BSA 647) and imaged for 8h. (**B**) Cells and grids images are segmented in binary images that are multiplied to isolate the grid nodes covered by the invasive boundary. Polygons tracking the spheroid spreading are reconstructed and averaged. (**C**) The time trend is projected and color-coded to visualize migration area A(t) (output #1). Diffusivity D(t) is obtained by differentiating A(t) (output #2). Spheroid boundary speed v(t) is obtained from D(t) and A(t)(output #3). Normalization of A(t) and v(t): τ, corresponding to the frame at which each cell line has migrated the same distance as the slowest cell line (C2) at the end of the acquisition (8h or more), and Δ, corresponding to the time window to complete 100 µm, are identified and all the movies are cut at τ. Running average (RA) movies are created by shifting Δ in the interval 1:τ and cell motility modes are characterized by extrapolating features from the RA movies: collective migration (output #4) is obtained by thresholding the area (outlined in green) of pixels belonging to the last bin of the histogram; directional persistence (output #5) is obtained by evaluating image orientation; hurdling (output #6) is obtained by sampling the intensities of the grid squares.

The second step of SP2G analysis provides the 3 other numerical outputs that characterize motility modes: collective migration (output #4: the higher the values, the more cells migrate as collective strands), directional persistence (output #5: the higher the values, the more cells move in the same direction) and hurdling (output #6: the higher the value the more cells are cutting angles). For the analysis of motility modes, only motile cells are considered. We defined motile cells by an average boundary speed higher than 100 μm / 8 h (0.21 μm/min). The characterization of motility modes is based on running average movies of the spreading spheroids and requires the identification of 2 parameters specific to each condition, Δ and τ, that allow normalization over migration area and boundary speed (see Supplementary File 1). Δ corresponds to the time window necessary to migrate 100 μm and τ corresponds to the frame at which each cell line has migrated the same distance as the slowest condition at the end of the acquisition (8h or more). Movies are cut at τ and running average (RA) movies are created by shifting Δ in the interval 1 to τ. Cell motility modes are characterized by extrapolating features from RA movies. Collective migration (output #4) is obtained by thresholding the area of pixels belonging to the last bin of the image histogram (outlined in green); directional persistence (output #5) is obtained by evaluating image orientation; hurdling (output #6) is obtained by sampling the intensities of the grid squares (i.e. the passivated areas covered by cells that cut angles) (Fig. 2C and S2). To test our SP2G analytical tool, we used simulated data of 100 round particles (radius=5 pixels) mimicking cells in spreading spheroid with different properties. Particles were diffusing at 3 speed regimes with full constraint (Continuous, 100% probability of being attached to a neighbor), partial constraint (Pseudo-continuous, 90% probability of being attached to a neighbor), or no constraint (Simple diffusion). As depicted in Fig. S2H, our analytical tool was able to distinguish accurately the different particle properties of collective migration.

### SP2G quantifies known migratory tactics

We tested our SP2G toolbox with 3 patient-derived cell lines known for their different motility modes on grids: NNI-11 (non-motile), NNI-21 (hurdler) and NNI-24 (glider) (*55*). As observed in fig. 3, SP2G confirmed their migratory behavior: within the same time window (4h), the most motile NNI-21 spread further away than the NNI-24 and NNI-11. This behavior could be quantified with diffusivity and boundary speed that were higher in the NNI-21 (Fig. 3A, F-G). The specific motility modes (hurdler vs glider) could also be quantified (Fig. 3D, H-K). Following our previous observations (*55*), NNI-21 cells migrated stochastically, with jumpy motions reflected by a low directional persistence (Fig. 3J) and high hurdling (Fig. 3K). Hurdling was visualized with cumulative distribution functions (CDF, lower slopes indicating more hurdling) and converted to numbers by dividing the average square intensity of the NNI-21 by the one of the NNI-24, returning a relative value of 2.08. NNI-24 displayed higher collective migration than NNI-21 (Fig. 3D, I ).

**Fig. 3.**
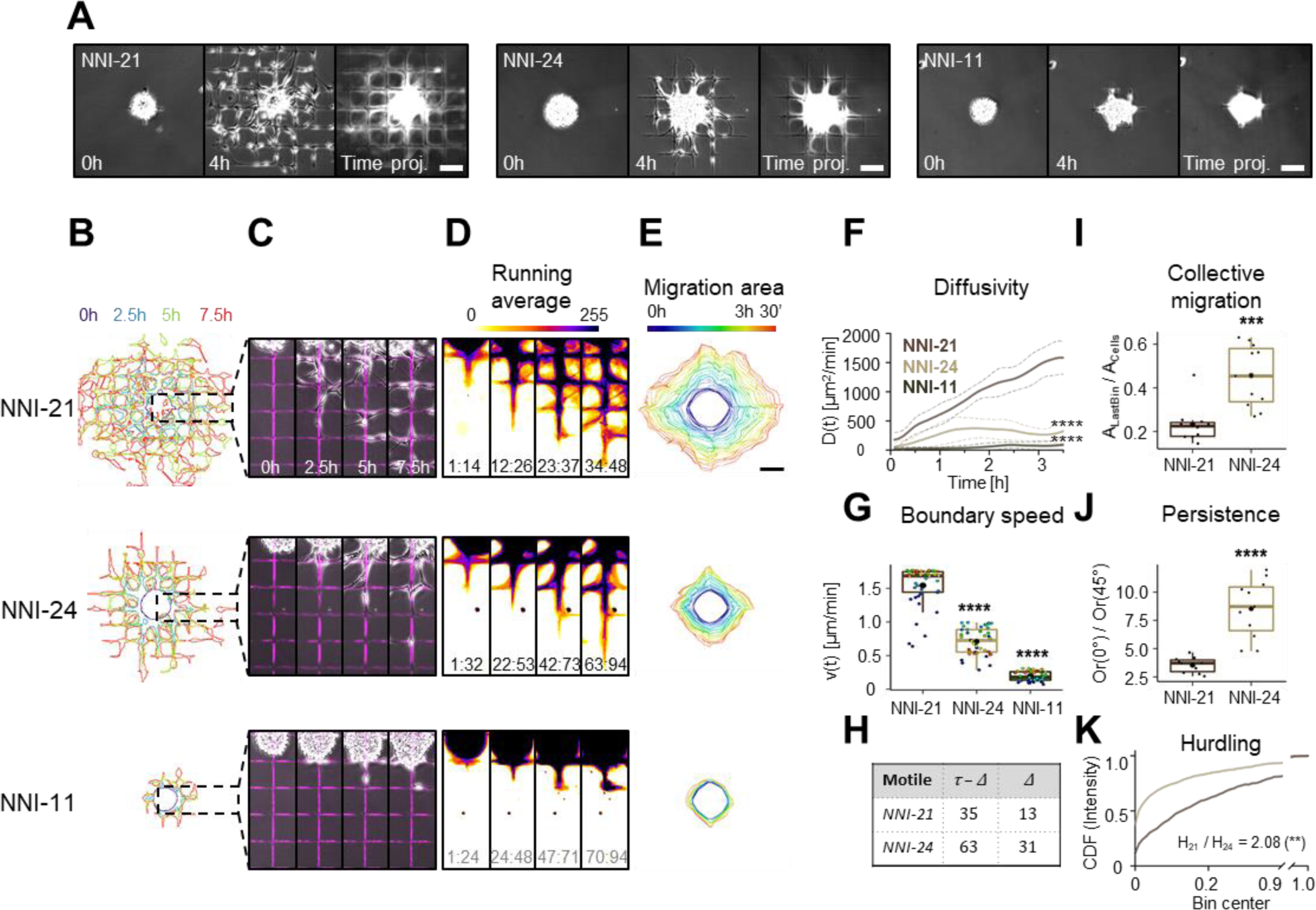
SP2G quantifies cell migratory tactics. Patient-derived glioblastoma spheroids were seeded on fluorescent gridded micropatterns, imaged for 8h, and analyzed as indicated in fig. 2 (n = 10, 11, 13 spheroids, n = 2, 2, 2 independent experiments for NNI-21, NNI-24 and NNI-11 respectively). (**A**) Snapshots of the movies at 0 h, 4 h, and corresponding time projections. (**B-D**) Cellular edges (**B**) and corresponding overlays of the phase contrast and the fluorescent grid images (**C**) at 4 time points (0h, 2.5h, 5h, 7.5h) and corresponding running average (RA) (**D**). The time window Δ constituting the corresponding RA frame is indicated at the bottom of each panel: for the non-motile τ = 94, Δ = 24. (**E**) Averaged polygons visualizing migration areas. (**F**) Diffusivity over 3.5 h; dashed lines are standard deviation. (**G**) Mean boundary speed over 3.5 h. Each dot represents a time-point and is color-coded as in (**E**). (**H**) Δ (number of frames needed to travel 100 μm) and τ - Δ (number of frames in the RA movie) in motile cell lines. (**I**) Collective migration quantification. Each dot represents a spheroid. (**J**) Directional persistence visualized as the ratio between the orientation along 0° and along 45° (Or(0°) / Or(45°), see methods). Each dot represents a spheroid. (**K**) Hurdling visualized as the Cumulative Distribution Function (CDF) of the normalized mean intensity of the grid squares (image intensity is sampled in each square). The ratio indicates the relationship between the average mean intensities (the sum of the mean intensity from all the squares divided by the total number of squares) of the 2 cell lines. Scale bars are 100 μm. Time and image intensity are color-coded as indicated. Statistical analyses are shown for NNI-21 compared to the other cell lines (see Supplemental for complete analysis).

We then evaluated SP2G sensitivity by applying a set of cytoskeleton-perturbing drugs to NNI-21 in a dose-dependent manner. We recorded the effects for the Arp2/3 inhibitor CK666, the myosin II inhibitor blebbistatin, the microtubule poison nocodazole, and the actin poison latrunculin-A (Fig. S3 and Supplementary Video 4). CK666 did not affect NNI-21 migration and motility mode, whereas all the other drugs reduced spheroid spreading and affected motility modes differently. Blebbistatin and latrunculin-A increased collective migration and persistence, and decreased hurdling while Nocodazole kept collective migration and directional persistence but slightly decreased hurdling. Overall, these results validated SP2G as a sensitive method for motility screening and highlighted its potency as a platform for drug testing.

Cells can sense changes in their microenvironment, including the chemical composition of the substrate. As observed previously, GBMs have a strong affinity for laminin (*53–56*). We tested SP2G sensitivity to perturbations of substrate density by micropatterning the grids with various laminin concentrations (400, 200, 100, 50, 25, 12.5, 6.25 μg/ml) and a blank condition (no laminin) (Fig. S4). Spheroids did not adhere in the blank and at the lowest concentration. Diffusivity and boundary speed measurements showed that cell migration increased with laminin concentration from 12.5 μg/ml up to 50 μg/ml where it reached a plateau (Fig. S4 E-F). Above 50 μg/ml LN, migration didn’t increase dramatically and didn’t decrease either. Interestingly, the boundary speed data indicated that the fastest cells were the cells monitored toward the end of the acquisition (red and yellow points) while the slowest were monitored at the beginning (dark blue points) (See Fig. 3G NNI-21, Fig. S3H control, Fig. S4F, LN 50-25-12.5) suggesting that cells accelerated with time after touching the substrate. However, at the highest LN concentrations (Fig. S4F, LN 200-400), the sorting of the cells was less obvious.

Altogether, SP2G emerged as sensitive in characterizing inter-patient heterogeneity in cell migration (NNI-11 vs 21 vs 24) and could detect subtle motility differences under fine biochemical perturbations (NNI-21). Therefore, we hypothesized that SP2G could unveil potential intra-patient cancer heterogeneity in migration and motility modes.

### SP2G reveals heterogeneity in the migratory tactics adopted by glioblastoma sub-populations isolated from patient-derived tumor-spheres

GBMs have been characterized both as inter- and intra-patient heterogeneous tumors. Heterogeneous GBM displayed difference in their genomic (*5, 11, 69*), epigenetic (*8, 70*), transcriptomic (*6, 7, 10, 12, 71*) and proteomic (*9*) profiles, which can be mutating under therapy (*72*) and maintained when cultured in vitro (*73*). Heterogeneity in motility has also been observed in mouse brains, but from these experiments it is unclear if the motile heterogeneity is due to the intrinsic properties of the cells or their surrounding environment (*74–76*). Using SP2G, we tested the motility heterogeneity of the patient-derived cell line GBM7 (Fig. 4 and Supplementary Video 5). Strikingly, when GBM7 spheroids spread on the grid, the single spheres separated in several small ones instead of spreading homogenously, suggesting that motile subpopulations were carrying non-motile cells, packed in mini spheroids (Fig. 4A and Supplementary Video 5). This implied that the GBM7 cell line was composed of heterogeneous populations displaying different intrinsic motility properties. Therefore, we decided to isolate these sub-populations and analyze their phenotype. Single clones were isolated by serial dilutions in 96 well plates and amplified as spheroids, from which we identified 3 motile (GBM7 clones #09, #01, #07) and 2 non-motile (#03, #02) sub-populations (Fig. 4B). The motile clones displayed two different cell morphologies. Clones #09 and #07 had small cell bodies and two thin polarized processes, whereas clone #01 displayed larger cell bodies, with multiple thick processes (Fig. 4B and Supplementary Video 5). From the live imaging, we could also observe that clone #01 was behaving differently than clones #07 and #09 (Supplementary Video 5). However, the 3 clones migrated with similar migration parameters (no significant differences in migration area and diffusivity with the exception of clone #09 that displayed higher boundary speed) (Fig. 4D-G, I-J). Clone #01 displayed higher collective migration, higher persistence and higher hurdling than clones #07 and #09 even though each of those differences were not significant (Fig. 4K-M). These results, together with the clear cell morphology differences observed on laminin, suggested that clone #01 belonged to another motile category than clone #07 and #09. The non-motile clones also displayed two different cell morphologies, with clone #03 presenting larger cell bodies and processes than clone #02. Both clones were not spreading on 2D or on grids, with low migration area, diffusivity and boundary speeds (Fig. 4D-G, I-J). To confirm SP2G as a bona-fide alternative to brain tissue assays, we tested these 5 populations in our brain slice assay. The brain slice assay confirmed the results obtained by SP2G (Fig. 4C,N and Supplementary Video 5), with 3 motile (#09, #01, #07) and 2 non-motile (#03 and #02) clones. However, a detailed quantitative characterization of behaviors and speeds was impossible with the brain slice assay. At the opposite, SP2G analysis, together with the cell morphology, could separate the 3 motile clones into 2 potential motile clusters: one cluster (clone #1) with higher hurdling, collective and persistence properties than the other cluster (clones #7 and #9) (Fig. 4B, K-M).

**Fig. 4.**
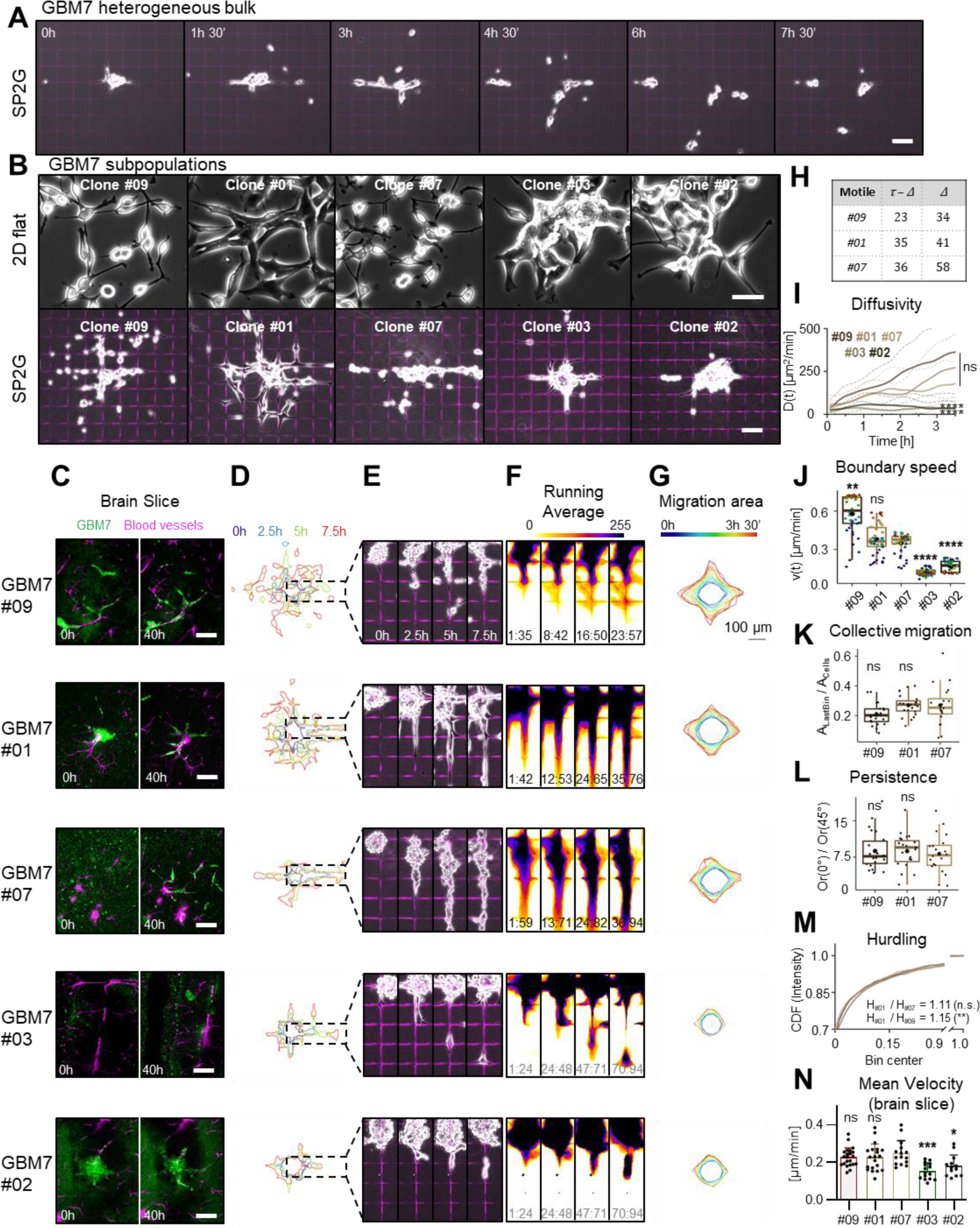
SP2G reveals migration heterogeneity in glioblastoma sub-populations isolated from patient-derived tumor-spheres. Spheroids from GBM7 original cell line (bulk) and isolated subpopulations (clones #01, #02, #03, #07, #09) were seeded on fluorescent gridded micropatterns, imaged for 8h, and analyzed as indicated in fig. 2. (**A**) Snapshots of the movies of the original population (bulk) at the indicated time-points. (**B**) Phase-contrast pictures of the GBM7 sub-populations cultured on laminin (top) and after SP2G at 8 hours. (**C**) Spheroid spreading of GBM7 sub-populations (green, DiOC6 dye) in brain slices at 0 h and 40 h. (**D-G**) SP2G analysis of the clones #01, #02, #03, #07, #09 (n = 22, 23, 20, 22, 22 spheroids respectively; n = 3 independent experiments): Cellular edges (**D**), corresponding overlays of the phase contrast and the fluorescent grid images at 4 time points (**E**), and corresponding running average (RA) (**F**). The time window Δ constituting the corresponding RA frame is indicated at the bottom of each panel: for the non-motile τ = 94, Δ = 24. (**G**) Averaged polygons visualizing migration areas. (**H**) Δ and τ - Δ of the motile subpopulations. (**I**) Diffusivity over 3 h 30’. Dashed lines are the standard deviation. (**J**) Mean boundary speed over 3h 30’. Each dot represents a time-point and is color-coded as in (**G**). (**K**) Collective migration for the motile cells. Each dot represents a spheroid. (**L**) Directional persistence for the motile cells visualized as the ratio between the orientation along 0° and along 45° (see methods). Each dot represents a spheroid. (**M**) Hurdling, visualized as the Cumulative Distribution Function (CDF) of the normalized mean intensity of the grid squares. The ratio indicates the relationship between the average mean intensities of the most hurdling (#01) against the others. (**N**) Mean velocities of single cells migrating in brain slices n = 20, 13 15, 15, 22 tracks for clones #1, 2, 3, 7, 9 respectively, each dot is a cell). Bars are 100 μm (**A, B bottom panel, C**) and 50 μm (**B, top panel**). Time and image intensity are color-coded as indicated. Statistical analyses are shown for clone #07 compared to all the other clones (see Data S3 for complete analysis).

In conclusion, these results confirmed that sub-populations hidden in patient-derived samples spanned a range of migration modes comparable to those from different patients and demonstrated that SP2G was a valuable platform to unveil them. Next, we asked whether the transcriptional profiles of our sub-populations could account for their differences in cell motility.

### Intra-patient heterogeneity in motility modes is correlated with specific molecular signatures

We profiled transcriptional landscapes of the clones GBM7 by RNA-seq to see if their signatures correlated with their motile phenotype defined by SP2G. RNA samples from three different cultures of each clone were sequenced by our onsite genomic unit (50.10^6^ reads per sample). Differential expression analysis followed by principal component analysis (PCA) showed that the motile (#01, #07 and #09) and the non-motile (#02, #03) clones grouped in 2 distinct clusters, and that inside the motile cluster, clone #07 and #09 grouped together away from clone #01, as suggested by SP2G and the morphology of the cells (Fig. 5A). Gene set enrichment analysis (GSEA) of differentially expressed genes in motile versus non-motile groups showed enrichment in the ECM-receptor interaction and focal adhesion pathways (KEGG) (Fig. 5B), EMT pathways (Hallmark), integrin and TGFβ pathways (Biocarta) (Fig. S5A-B) that are all linked to motility. Strikingly, z-score of expression levels of integrin genes indicated that the laminin-binding integrins (particularly ITGA3 and ITGA6) were enriched in the motile clones compared to the non-motile (Fig. 5C). In opposition, fibronectin-binding integrins were either poorly expressed or uncorrelated to cell motility (Fig. 5C). We confirmed these results by qPCR that showed that the laminin-binding integrins ITGA3 and ITGA6 were enriched in the motile clones while the fibronectin-binding integrin ITGAV was not (Fig. 5E). ITGA5, ITGA7 and ITGA10 were also enriched but to a lower extent. These results were confirmed by western blot (Fig. 5D) and are in agreement with glioma preference for laminin and with the large presence of laminin on brain blood vessels (*53–56, 62–67*). The comparison of the top 50 upregulated genes in the motile group versus the non-motile (Fig. S5C) showed that while most of the genes that were upregulated in clones #07 and #09 were also upregulated in the clone #01, some genes such as LMO3 and HHIP were upregulated only in clone #01 and others such as CRABP2, MOXD1, MTL5, CTSZ were upregulated in clones #07 and #09 but not in clone #01 confirming the clustering of the motile clones in 2 different groups. On the other hand, all the top 50 downregulated genes showed the same trends in all the 3 motile clones (Fig. S5D).

**Fig. 5.**
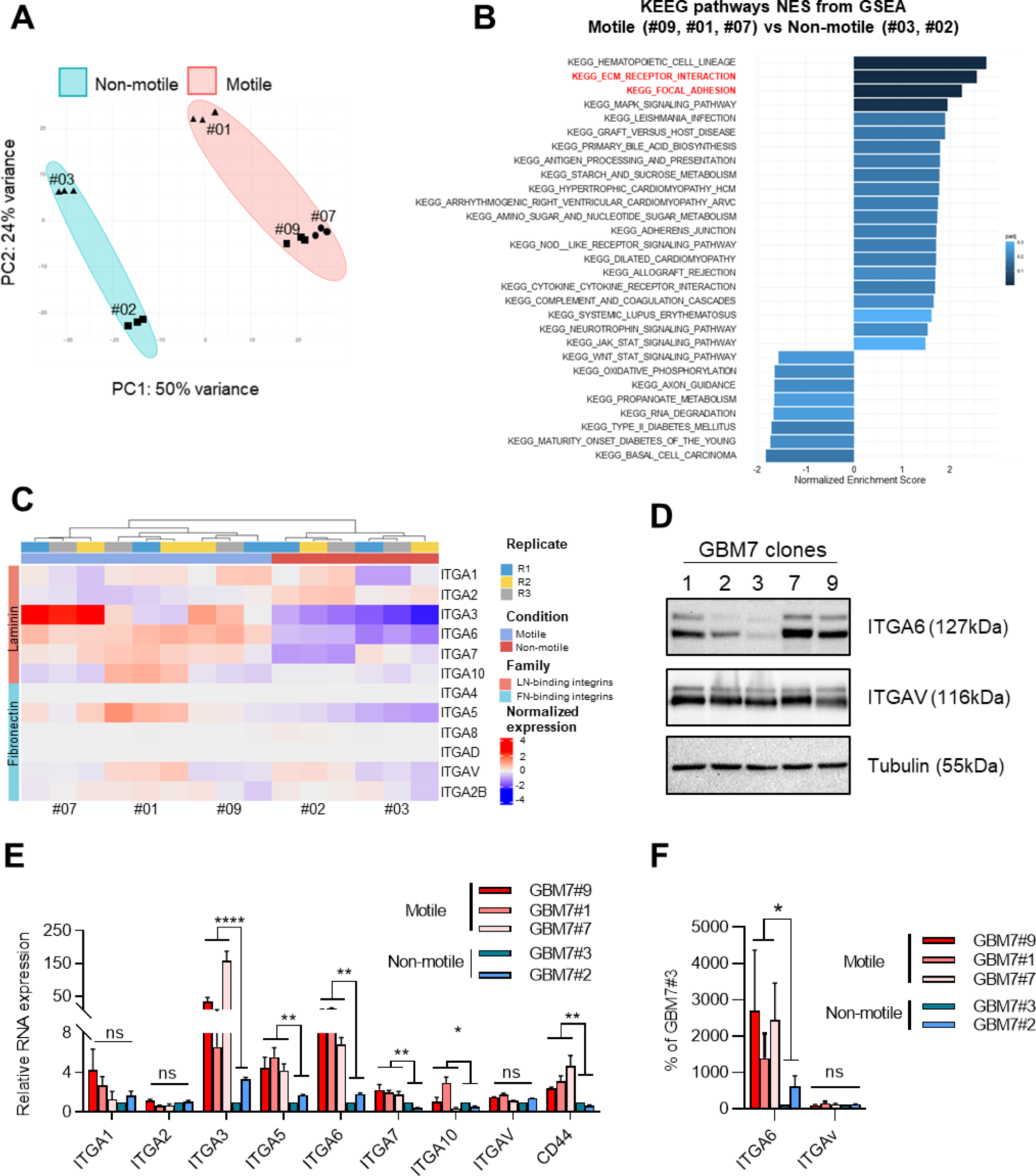
Intra-patient heterogeneity in migration ability is correlated with specific molecular signatures. (**A**) Principal component analysis showing segregation of the 5 GBM7 sub-populations in motile and non-motile groups. (**B**) Gene set enrichment analysis (GSEA) of differentially expressed genes in the motile vs non-motile group. GSEA was performed using the Kyoto Encyclopedia of Genes and Genomes (KEGG) gene set in the GSEA molecular signatures database. Moderated t-statistic was used to rank the genes. Reported are Normalized Enrichment Scores (NES) of enriched pathways (with the fill color of the bar corresponding to the P-value). P-value was calculated as the number of random genes with the same or more extreme ES value divided by the total number of generated gene sets. (**C**) Heatmap representing z-score of expression levels of integrins. (**D**) Expression of ITGA6, ITGAV, tubulin, in total cell extracts of the 5 GBM7 sub-populations growing on laminin. (**E**) Relative mRNA expression levels of ITGA1, ITGA2, ITGA3, ITGA5, ITGA6, ITGA7, ITGA10, ITGAV, and CD44 in the 5 GBM7 sub-populations. Each integrin expression is reported to its expression in clone #3. GAPDH and B2M were used as housekeeping genes. n=3 independent experiments. Error bars are S.E.M. (**F**) Quantification of the expression of ITGA6 and ITGAV in each condition reported to their expression in clone #3. 2 independent western blots were quantified.

To assert more precisely the differences between the 2 motile groups, we analyzed them without the non-motile. The differential expression analysis followed by principal component analysis (PCA) showed 75% variance between the clone #01 and the 2 remaining clones while the variance between the clone #07 and #09 was only 19% (Fig. 6A). Interestingly, Gene set enrichment analysis (GSEA) of differentially expressed genes showed a depletion in the EMT pathway (hallmark), and in the ECM-Receptor interaction and focal adhesion (KEGG) suggesting that the clone #01 was more mesenchymal than clones #07 and #09 (Fig. 6C, S6A). This was confirmed with the comparison of the top variable 20 genes between clone #01 and clones #07-09 that contained genes involved in the EMT and matrix organization with TGFβ1 being the most variable gene between these clones (Fig. 6B). Within these twenty genes, 6 encoded for extra cellular matrix proteins: EMILIN1 (Elastin Microfibril Interfacer 1, that is involved in Elastic fiber formation and Extracellular matrix organization), MGP (Matrix Gla Protein, that is normally secreted by chondrocytes and vascular smooth muscle cells, and functions as a physiological inhibitor of ectopic tissue calcification); XYLT1 (Xylosyltransferase 1, necessary for biosynthesis of glycosaminoglycan chains), FN1 (Fibronectin), COL4A5 (Collagen Type IV Alpha 5), NCAN (Neurocan, a chondroitin sulfate proteoglycan thought to be involved in the modulation of cell adhesion and migration). Hence, we could assign different transcriptional signatures to the two motile behaviors. Taken together, these results validated the motile versus non-motile classification as well as subtle distinctions in motility mode, and provided insights on the molecular determinants that characterize GBM intra-patient heterogeneity in cell motility.

**Fig. 6.**
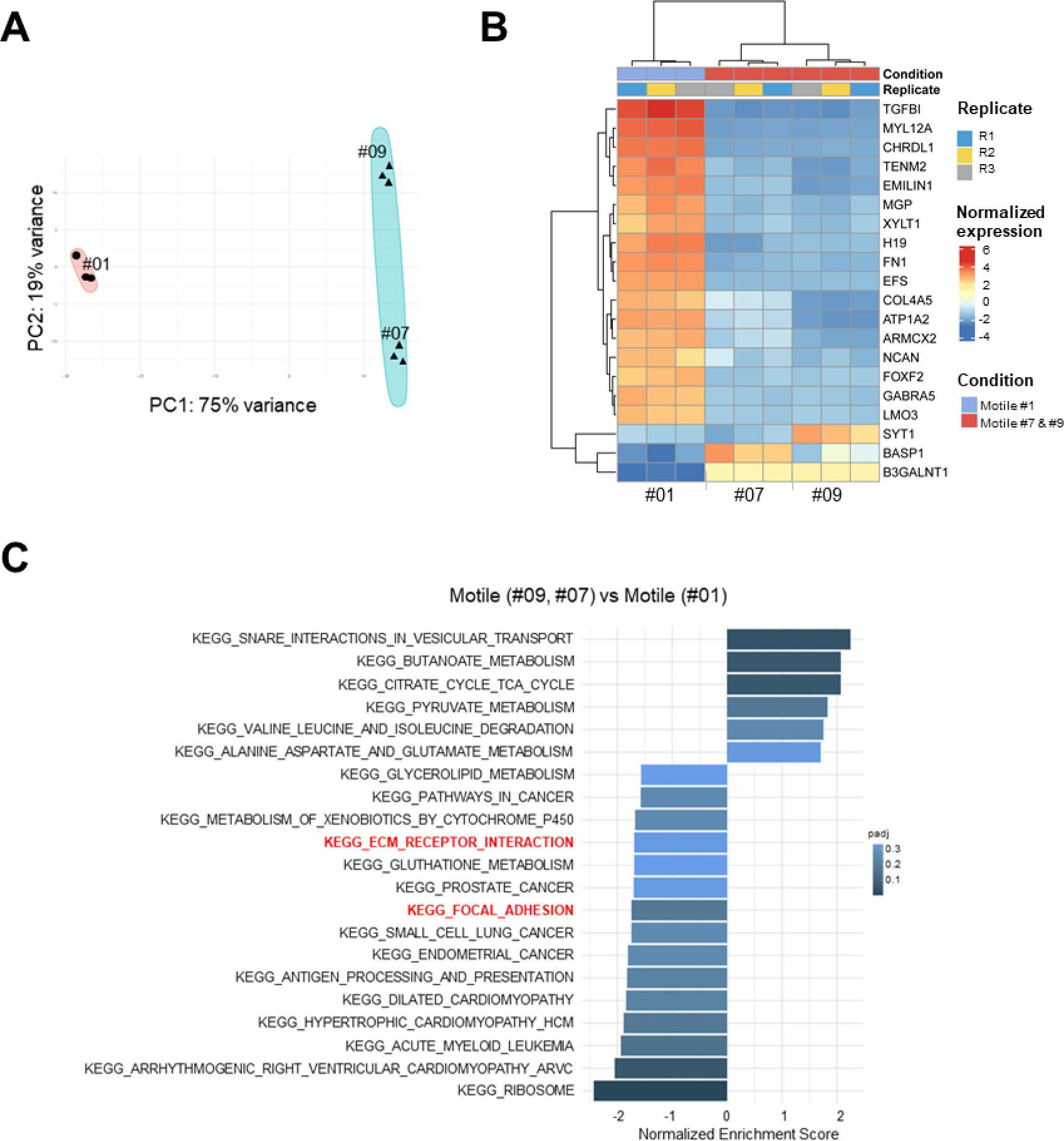
Intra-patient heterogeneity in motility modes is correlated with specific molecular signatures. (**A**) Principal component analysis showing segregation of the 3 motile sub-populations in 2 groups. (**B**) Heatmap representing row-normalized expression levels of the 20 most variable genes between the 2 motile groups. Genes belonging to the EMT pathway are upregulated in clone #01 compared to clones #07 and #09. (**C**) Gene set enrichment analysis (GSEA) of differentially expressed genes in the motile groups (clones #07 and #09 versus clone #01). GSEA was performed using the KEGG gene set in the GSEA molecular signatures database as in fig. 5.

## DISCUSSION

In heterogeneous tumors such as glioblastoma, some cells can be highly aggressive, migrating long distances from the tumor core while other cells can be less motile, remaining in the tumor core (*7, 71, 74–78*). Within the motile populations, cells may also display significant heterogeneity in term of motility modes and molecular signatures, and in consequence display different sensitivity to targeting drugs. Our goal, here, was to develop a method to analyze rapidly the various motility modes present in single patients in order to identify the most aggressive clones and their specific molecular signatures. We tested SP2G using patient-derived cell lines NNI-11, NNI-21, and NNI-24, which have known migration behaviors. In our previous work, we extensively studied the migration and motility modes of these cell lines (*55*). NNI-11 showed non-motile and highly proliferative behavior, while NNI-21 and NNI-24 exhibited motile characteristics with distinct patterns: NNI-21 showing stochastic, jumpy motion (hurdlers), and NNI-24 following defined tracks (gliders). SP2G outperformed our previous system with many advantages:

1. SP2G allowed a faster, semi-automated, analysis of the cell migration, by tracking the spreading area of the spheres instead of manually tracking single cells like in (*55*). The differences in motility were clearer and more complete than our previous results. NNI-11 were not spreading at all, and NNI-21 and NNI-24 mirrored their known migration proficiency (for NNI-21: 90 μm/h with SP2G vs 60 μm/h for single cells on grid; for NNI-24: 40 μm/h with SP2G and 30 μm/h single cell on grid (*55*).
2. SP2G could quantify the hurdling of NNI-21 as opposed to the high directional persistence of NNI-24. Originally, we defined hurdling and gliding with the projection of all the frames of a 6h movie. The projection of glider movies resulted in a quasi-perfect grid, drawn by the cells, while the projection of hurdler movies resulted in grids that were filled-up in the non-adhesive area with cell processes (*55*). However, quantification was not straightforward and could depend on the cell density. With SP2G, hurdling evolution could be analyzed in time and cell lines could be compared. Hence, SP2G can define and score hurdling motility in an unbiased way. Using different doses of cytoskeleton-perturbing drugs, we demonstrated that SP2G was sensitive and suitable as a motility-screening platform. We confirmed the requirement of myosin II and microtubules and the independence of Arp2/3 for the motility of GBM cells as we previously reported (*55, 56*).
3. SP2G highlighted the formation of collective strands as observed in vivo (*22, 24, 46*), that cannot be recapitulated using the single cell migration assay. Indeed, the spheroid acts as a reservoir for the cells to spread diffusively, allowing many cells to move in the same direction. Moreover, the short duration of our experiments (8 hours) reasonably ensures the independence of cell migration from cell proliferation. SP2G highlighted and quantified the formation of collective strands in the NNI-24. These collective migration properties could not be identified in our previous work with single cell assays (*55*).
4. SP2G allows the migration of cells on a highly controlled substrate, in shape and in composition that has not been in contact with cells. In this regards, SP2G mimics the conditions encountered by the first migrating cells leaving the tumor core and invading the naïve blood vessel surface. SP2G can be adapted to any matrix proteins, allowing the analysis of other cell type than glioma. By modifying the concentration of laminin of the grid, we found that the NNI-21 were sensitive to small substrate variations and SP2G was able to pick these subtle differences. Interestingly, diffusivity and boundary speed measurements showed that cell migration increased with laminin concentration up to 50 µg/ml where it reached a plateau above which migration didn’t change. This migration increased with laminin concentration was previously observed with the GBM cell lines GaMG and U373 (*66*). These results are in opposition with previous modelling and experimental studies with different cell types, showing that cell migration exhibited a biphasic response to changes in fibronectin concentration (*79, 80*) or in laminin concentration for melanoma cells, revealing a dose optimum of 10 µg/ml laminin (*66*). Our results suggest that the laminin –binding integrins involved in GBM cells are behaving differently than the fibronectin-binding integrins, leading to variations in cell migration. Moreover, the boundary speed data indicated that cells accelerated with time after touching the substrate. This could be due to a gradually increase in the expression of pro-migration molecules triggered by the laminin substrate. In accordance, we previously showed that the expression of the formin FMN1 increased upon increased laminin concentrations in NNI-21 (*55*). At the highest concentration, however, the sorting of the cells was less obvious, suggesting that at high LN concentration cells expressed pro-migratory factors at earlier time points.
5. SP2G can unveil migratory heterogeneity and render it easy to see by eye. This is an important feature of SP2G. Identifying and demonstrating cell heterogeneity with single cells seeding on the grid would necessitate single cell tracking coupled to a deep analysis of the shape of each cell. Moreover, cells that do not adhere very well on the grid and are floating around could be under-evaluated. Here, by seeding spheroids of the bulk GBM7 cell line, we could detect the presence of migration heterogeneity right away, because single spheres separated in several small ones instead of spreading homogenously, suggesting that motile subpopulations were carrying non-motile cells, packed in mini spheroids. We do not know if this behavior occurs *in vivo* which would drive non-motile GBM cells away from the tumor core. However, we know that such ‘hitchhiking mechanisms’ exist for other cancer cells like melanoma (*81*). In this regard, this phenomenon could be tested via SP2G with mixed spheroids containing motile and non-motile populations. Correlative single cell RNA-seq in patient samples could then help deciphering if indeed motile cells could carry non-motile cells away from the tumor core *in vivo*. If this is the case and that non-motile cells can be found at the tumor margin, then, previous statements using high throughput approaches comparing tumor core and tumor margin, without corresponding cell migration assays might need to be re-evaluated (*82*). Also, motile and non-motile cells might communicate via networks or extra cellular messengers. Our method could be adapted to analyze the influence of the clones on each other, as it has been suggested (*45, 75, 83*).
6. SP2G can pick subtle differences and translate ‘eye’ observations into numbers. When we analyzed the GBM7 clones, we could observe that the motile clone GBM7 #01 was behaving differently than the motile clones #07 and #09. Importantly, these motility clusters were assembled similarly using RNA-seq. In accordance with their motility on laminin, the 3 motile clones differed from the non-motile clones with the high expression of the laminin-binding integrins ITGA3 and ITGA6 while there was no difference for the fibronectin-binding integrin ITGAV. This also confirmed that the motility on laminin reflected better what it is happening in the brain since our motile and non-motile clones on laminin were the same motile and non-motile on brain slices. The fact that they did not displayed differences in the expression of fibronectin-binding integrin ITGAV could explain the negative results obtained with cilengitide, the integrin alpha V inhibitor, in clinical trials (*15, 16*): these cells may not use fibronectin-binding integrins to invade the brain. The expression of ITGA6 in the motile GBM7 clones could also reflect on their stemness, since integrin alpha 6 is known to regulate glioblastoma stem cells (*84*). At this stage, we do not know whether there is a correlation between stemness markers and sub-population motility. The integration of SP2G data and RNA-seq analysis of the clones, combined with observations on 2D dishes, revealed the existence of two distinct motile behaviors. Through the corresponding RNA-seq data, we can now outline a molecular signature that sets apart these two motile behaviors, with a notable EMT signature predominantly present in clone #01.

So far, GBM heterogeneity was mostly studied with genomics and transcriptomic tools (*5–8, 10–12, 69–71, 73, 77, 85, 86*). Here, we are demonstrating intra-patient heterogeneity at the phenotypic level by reporting how 5 sub-populations isolated from a single patient-derived sample migrate differently. Our RNA-seq data implied that specific transcriptional signatures could define motility modes and that the motile behavior is not driven only by the environment (*73, 76, 87, 88*) but some of its core component are anchored within the cell. Interestingly, differential transcriptional signatures have been linked to different invaded brain locations in xenograft experiments (*17, 87*). These preferential localizations maybe due to the various motility modes that various cell lines are using and it would be interesting to connect motility modes defined by SP2G with brain localization and transcriptional signatures.

Overall, these results highlight SP2G strengths in identifying motility modes with great details and a level of refinements hard to reach with other experimental approaches.

In summary, we have presented a methodology that integrates the time-lapse imaging of spheroid spreading on grids with an ImageJ/Fiji analytical toolbox that quantitatively characterizes cell migration and motility modes. It is nicknamed SP2G, and we hope it opens up a new standard for motility screenings, potentially extendable as a pan-spheroid approach that helps answering questions on how cell migration affects cancer dissemination.

## Materials and Methods

### Cell culture

Rat C6 cells were cultured in high-glucose DMEM supplemented with glutamine and 10% fetal bovine serum (FBS). To form spheroids, ∼2·10^6^ C6 cells were seeded in 6-cm petri dishes previously treated for 1h with 0.2% pluronic F127 at room temperature. After 1 day, spheroids between 75 and 150 μm in diameter were obtained. Patient-derived GBM cell lines from the laboratory of Carol Tang at the National Neuroscience Institute in Singapore (NNI-11, NNI-21 and NNI-24) were collected with informed consent and de-identified in accordance with the SingHealth Centralised Institutional Review Board A. Patient-derived cell line GBM7 from the laboratory of G. Pelicci (IEO, Milan, Italy) was collected according to protocols approved by the Institute Ethical Committee for animal use and in accordance with the Italian laws (D.L.vo 116/92 and following additions), which enforce EU 86/609 Directive (Council Directive 86/609/EEC of 24 November 1986 on the approximation of laws, regulations and administrative provisions of the Member States regarding the protection of animals used for experimental and other scientific purposes). All patient-derived GBM cell lines were kept as previously reported (*89*). Briefly, GBM cell lines were grown in non-adherent conditions in DMEM/F-12 supplemented with sodium pyruvate, non-essential amino acid, glutamine, penicillin/streptomycin, B27 supplement, bFGF (20 ng/ml), EGF (20 ng/ml), and heparin (5 mg/ml). Patient-derived GBM cell lines were passaged every 5 days. All the cell lines were maintained at 37 °C and 5% CO2.

### Brain slice invasion assays and staining

C57BL/6J mice were used for these studies. Both males and females (in equal proportions) within each experiment originated from different litters. All of the animal procedures were in accordance with the Institutional Animal Care and Use Committee, and in compliance with the guidelines established in the Principles of Laboratory Animal Care (directive 86/609/EEC); they were also approved by the Italian Ministry of Health. The brain slice assay was performed as reported in Er et al. (*90*) and Polleux and Ghosh (*91*). Prior to sacrifice, mice were anesthetized, their chest was cut and intra-cardiac injection was performed with 5 ml solution of Dil stain to label the luminal side of blood vessels. The Dil stain was diluted at 0.5 mg/ml in 100% ethanol and this solution was further diluted 1:10 in a 30% w/v solution of sucrose-DPBS. Brains were then isolated in ice-cold CaCl2+/MgCl2+ 1X HBSS (Euroclone ECB4006) supplemented with 2.5mM HEPES (complete HBSS). Brains were sectioned in 150 or 100 μm thick slices using a Leica VT1200S vibratome and placed in a glass bottom 24-well, which was previously coated at 37 °C overnight using a solution of 12.5 mg/ml laminin and 12.5 mg/ml Poly-L-lysine (1 slice/well). Slices were left 3h at 37 °C and 5% CO2 to consolidate on the substrate. Subsequently, glioma spheroids were gently added and the co-culture was kept 4h at 37 °C and 5% CO2 prior to imaging. Movies were recorded for 72h on a confocal SP5 microscope equipped with temperature, humidity, and CO2 control utilizing a 20X air objective (1 frame/15 min for rat C6, 1 frame/30 min for GBM-7 sub-populations). All the brain-slice live experiments were performed with brain-slice culture medium (68% L-glutamine supplemented DMEM, 26% complete HBSS, 5% FBS, 1% Penicillin-Streptomycin). For immunofluorescence staining of C6 and blood vessels, the co-culture was fixed with 4% PFA for 20 min and incubated for 1h at room temperature with a blocking solution (5% BSA, 5% Normal-Donkey-Serum (NDS), 0.3% Triton-X 100 in DPBS). The co-culture was incubated overnight at 4 °C with 10 µg/ml of Tomato Lectin (Vector Laboratories, #DL-1178) in blocking solution. DAPI was added afterwards. Images were acquired with a Leica SP8 microscope utilizing a 63X oil objective (1 μm Z step).

### Collagen and reconstituted basement membrane (rBM) invasion assays

Collagen and rBM assays were adapted from Shin et al., (*92*). Briefly, 20 ml of polydimethylsiloxane (PDMS; Sylgard 184 Dow Corning) were casted at 1:10 ratio by mixing curing agent and silicone elastomer base, respectively, in a 10-cm plate. 6-mm PDMS wells were obtained by punching holes with a biopsy puncher in 18×18 mm PDMS squares. The PDMS wells were bound on 24 mm coverslips via plasma treatment (90 s) followed by 5 min at 80 °C. Each 6-mm well was then coated with 1 mg/ml poly-d-lysine at 37 °C for 3h, rinsed in milliQ water, and cured overnight at 80 °C. Meanwhile, rat C6 spheroids were incubated in medium with 5 μM Dil stain for 3 h, centrifuged at 500 rpm for 2 minutes and re-suspended in 1 ml of medium. 10 μl of spheroid suspension was then mixed with 80 μl of 6 mg/ml collagen solution (Collagen I diluted in cell culture medium, 10% v/v 1.2% NaHCO3, 5% 1M HEPES, 1.5% 1M NaOH) or 10 mg/ml reconstituted basement membrane (rBM). The spheroids embedded in unpolymerized solutions were placed in the 6-mm PDMS wells and left at 37 °C for 1h to polymerize. With this method 5 to 15 spheroids per well were obtained. Medium was then added and movies of invading spheroids were acquired on an IX83 inverted microscope (Olympus) equipped with a Confocal Spinning Disk unit, temperature, humidity, and CO2 control. Images were collected with a 10X objective, an IXON 897 Ultra camera (Andor) and OLYMPUS cellSens Dimension software for > 24 hours (1 frame / 15 min) with 7.5mm Z step for RFP and DIC channels.

### Quantification of spheroid spreading

Spheroid spreading quantification in collagen and reconstituted basement membrane (rBM) assays was obtained as the ratio between the area occupied by spheroids at 0 and 24 hours. The area was calculated as the maximum intensity projection of the fluorescent channel in Z and in time. For spheroid spreading quantification in 2D flat and grid assays, the ratio between the areas at 8 and 0 hours was calculated, and the areas were obtained using the maximum intensity projection in time. For MSD calculation we applied the protocol of Gorelik at al. (*93*). XY coordinates over-time were obtained by manual tracking with the dedicated plugin in Fiji. For brain slice and 3D gels assays, cells were tracked for 14 h. To track well-identified single cells, initial time points were taken several hours after the start of the acquisition (24 h for brain slice and 10 h for 3D gels).

### Microcontact printing

Microcontact printing was performed as we previously described (*50, 89*). Briefly, we casted 1:10 PDMS from a dedicated silicon mold, cut it into 1×1 or 1×2 cm^2^ stamps, and coated with 50 μg/ml laminin in DPBS for 20 min. Each stamp was then air-blow dried, leant on a 35-mm dish, then gently removed. In case of plastic dishes, the surface was passivated with 0.2% pluronic F127 in DPBS at room temperature for 1 h, whereas for glass poly-l-lysine-grafted polyethylene glycol (0.1 mg/ml, pLL-g-PEG, SuSoS) was used. Dishes were then rinsed 4 times with DPBS and kept in medium until spheroids were seeded. For printing laminin concentrations from 400 to 6.25 μg/ml, a sequence of 6 serial dilutions (1:1 in DPBS) was carried out.

### SP2G experimental and image analysis workflow

To stain the micropatterns, the laminin solution for was mixed with 7 μg/ml BSA-conjugated-647. To stain the spheroids, 1-day old (for rat C6 glioma cells ) or 5-days old spheroids (for human patient-derived glioma) were incubated in medium with 5 μM Dil stain for 3 hours in 6-well plates previously passivated with 0.2% pluronic F127. Spheroids were then seeded on the grid and the samples were placed under the microscope and left 5 to 15 min to equilibrate. 8-hours time-lapse movies were recorded using a 10X objective mounted on a Leica AM TIRF MC system or onto an Olympus ScanR inverted microscope (1 frame / 5 min). 3 channels per time point (phase contrast, Dil stain fluorescence for the cells, BSA-647 fluorescence for the grid) were acquired in live cell imaging for the experiments in Fig. 3 and Fig. S4. 2 channels per time point were acquired in the other experiments, since the grid fluorescence was recorded for just 1 frame before and 1 frame after the time-lapse movie. For the Experiments in Fig. S4 cells were not fluorescently labeled, the drugs were injected 25 min after starting the acquisition. For the characterization of cell migration, we used the SP2G analytical toolbox that measured the polygonal area A(t). Briefly, as SP2G formed the polygon to track the invasive boundary, the code initially multiplied the binarized grid nodes with the binarized spreading spheroid in order to obtain only the node traveled by at least 1 cell. Then, SP2G iteratively checked whether a node has blinked for at least 3 consecutive time frames to add the node to the polygon. Therefore, SP2G always stopped tracking 2 frames earlier than the total duration of any movie.

From A(t) we derived the diffusivity D(t) = dA(t) / dt, where dt was the time frame in the time-lapse movies (5 min) and dA(t) was the difference between 2 polygonal areas at subsequent time steps. For the calculation of single cell velocity 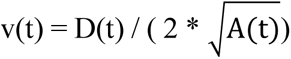, being now 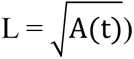 the edge of the square having an area equivalent to the polygon, the following 2-equation system was solved:

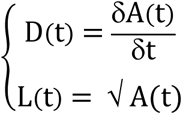

That inferred

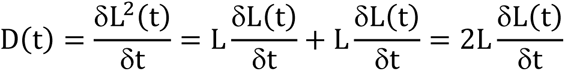

The following was obtained

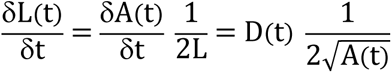

that corresponded to the value of the boundary speed. In this way, a length gradient was inferred from an area gradient.

For the characterization of the motility modes, we extrapolated all the parameters from RA movies and averaged data from several spheroids. Collective migration values were obtained by thresholding each frame of the RA within the last histogram bin, that necessarily spans up to 255 (the maximum value of an 8-bit image). The ratio

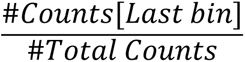

returns the collective migration. The rationale behind this assumption was that collective strands generated high intensity values when averaged, since many cells travelled the same path. Therefore, in the RA movie there were zones of high intensity. Vice versa, single entities generated low intensities when averaged, since no cells other than the single one contributed to the final average value. Background pixels were set to NaN (see Supplementary Appendix).

Directional persistence is calculated through the function “OrientationJ distribution” of the OrientationJ plugin (*94*), which returns the orientation field (OF): it consists in 180 values (1 per direction, sampled every 1°). Reasonably, we assumed that the spheroid spreading is isotropic, and therefore SP2G averages the values 0-90, 1-91, etc. and gets 90 values. The following ratio

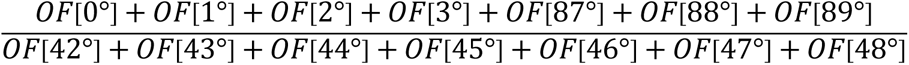

returns the directional persistence. It is the ratio between the direction of least resistance to cell migration (i.e. the ones provided by the grid segments) and the direction of most resistance (the one a cell has necessarily to face when undergoing a directional change).

### Simulation of particle diffusion

Simulated data were generated with a custom-written code in imageJ/Fiji. Briefly, the function

Speed_particle = speed*(1+random(“gaussian”));

was applied at each time step to generate motion. “speed” was equal to 2, 2.5 or 3 and “random(“gaussian”)” returned a Gaussian distributed pseudorandom number with mean 0 and standard deviation 1. Continuity was set by imposing a 100% overlap probability to moving particles, pseudo-continuity a 99% probability, pure diffusivity with no constraints.

### RNA extraction and qPCR analysis

Cells cultured on laminin were lysed and RNA was extracted with the RNAeasy Mini Kit (Qiagen) as per manufacturer’ specifications. About 1 µg of RNA was retrotranscribed using ‘qScript cDNA synthesis kit’ (Quantabio). For gene expression analysis, 5 ng of cDNA was amplified (in triplicate) in a reaction volume of 10 µl containing 5 µl of TaqMan® Fast Advance Master Mix and 0.5 µl of TaqMan gene expression assay 20X (Thermofisher). The entire process (retrotranscription, gene expression, and data analysis) was performed by the qPCR service at Cogentech-Milano, following ABI assay ID data base (Thermo Fisher). The qPCR sets of primers for ITGA1 (Hs00235006_m1), ITGA2 (Hs0018127_m1), ITGA3 (Hs01076873_m1), ITGA5 (Hs00233732_m1), ITGA6 (Hs00173952_m1), ITGA7 (Hs01056475_m1), ITGA10 (Hs01006910_m1), ITGAV (Hs00233808_m1), CD44 (Hs00153304_m1) and the housekeeping genes GAPDH (Hs99999905_m1) and GusB (Hs99999908_m1) were from Thermofisher. Real time PCR were carried out on the 7500 Real-Time PCR System (Thermo Fisher), using pre-PCR step of 20 s at 95 °C, followed by 40 cycles of 1 s at 95 °C and 20 s at 60 °C. Samples were amplified with primers and probes for each target, and for all the targets, one NTC sample was run. Raw data (Ct) were analyzed with Excel using the ΔΔCT method to calculate the relative fold gene expression. ΔCT was calculated using 2 housekeeping genes and averaged (3 independent experiments). For the mRNA expression of selected integrins, data were normalized against the expression of the GBM7 sub-population #03.

### RNA-sequencing and analysis

RNA-sequencing was performed by the qPCR service at Cogentech-Milano. Prior sequencing, RNA concentrations were measured using Qubit 4.0 and RNA integrity evaluated with an Agilent Bioanalyzer 2100 utilizing Nano RNA kit (RIN > 8). An indexed-fragment library per sample was arranged from 500 ng total RNA using Illumina Stranded mRNA Prep ligation kit (Illumina) as per manufacturer’s instructions. Libraries were checked for proper size using Agilent Bioanalyzer 2100 High Sensitivity DNA kit, then normalized and equimolarly pooled to perform a multiplexed sequencing run and quantified with Qubit HS DNA kit. As a positive control, 5% of Illumina pre-synthesized PhiX library was incorporated in the sequencing mix. Sequencing was carried out in Paired End mode (2×75nt) with an Illumina NextSeq550Dx, generating on average 60 million PE reads per library. Reads were aligned to the GRCh38/hg38 assembly human reference genome using the STAR aligner (v 2.6.1d) (*95*) and reads were quantified using Salmon (v1.4.0) (*96*). Differential gene expression analysis was performed using the Bioconductor package DESeq2 (v 1.30.0) (*97*) that estimates variance-mean dependence in count data from high-throughput sequencing data and tests for differential expression exploiting a negative binomial distribution-based model. The Bioconductor package fgsea (v 1.16.0) (*98*) and GSEA software (including Reactome, KEGG, oncogenic signature and ontology gene sets available from the GSEA Molecular Signatures Database, https://www.gsea-msigdb.org/gsea/msigdb/genesets.jsp?collections) were used for preranked gene set enrichment analysis (GSEA) to assess pathway enrichment in transcriptional data.

### Protein extraction and western blots

Total cell extracts were prepared in RIPA buffer (100 mM NaCl; 1 mM EGTA; 50 mM Tris pH7.4; 1% TX100) complemented with a cocktail of protease inhibitors (Roche). Proteins were quantified using the Pierce BCA protein assay kit #23225 (ThermoScientific). Proteins were denatured with SDS and resolved by SDS-PAGE using typically 8% acrylamide gels. Transfers were done on PVDF membranes in methanol-containing transfer buffer. Blocking was done with milk diluted to 5% in PBS-0.1% tween for 1h at room temperature and antibodies were blotted overnight at 4 °C. HRP-secondary antibodies were incubated for 1-2 hours at room temperature in 5% milk and ECL were performed using the Amersham ECL Western Blotting Detection Reagents (Cat.no. RPN2106 from GE Healthcare). Detection was done using the Chemidoc XRS imaging system (Bio-Rad)..

### Reagents

Rabbit anti-integrin alpha V antibody (ab179475) was from abcam. Rabbit anti-integrin alpha 6 (NBP1-85747) was from Novus. Mouse Anti-tubulin (T9026) antibody was from Sigma. Laminin (#23017015), Dil stain (#D282) and BSA-conjugated-647 (#A34785) were from ThermoFisherScientific. Collagen I was from Corning (#354249). Reconstituted basement membrane (rBM) was from Trevigen (# 3445-005-01).

### Statistical analysis

All the statistical analysis was performed with Prism 9 (GraphPad). Number of samples and independent experiments are indicated in figure legends. The plots were generated with Prism 9 and ggplot2. In all the boxplots, the middle horizontal line represents the median and the black dot is the mean value. *** means p<0.001, **** means p<0.0001. All the statistical analyses are detailed in the Excel file ‘data S3’. Radar plots were generated with Google sheets.

### Codes and macros

All the SP2G macros and the supplementary appendix containing detailed instructions on installation and run are freely available on the repository figshare at the following link: https://figshare.com/projects/SP2G/148246.

## Acknowledgments

We are grateful to the IFOM imaging facility personnel, in particular D. Parazzoli for technical support, the IFOM cell culture facility personnel, the MBI microfabrication facility personnel, in particular Sree Vaishnavi Sundararajan. We thank Marco Foiani and Giorgio Scita (IFOM), Virgile Viasnoff (MBI), Marc-Antoine Fardin (Institut Jacques Monod), Nir Gov (Weizmann Institute of Science, Israel), Scita’s and Maiuri’s groups for helpful discussions and critical comments on the manuscript.

## Funding

This work was supported by:

IFOM (starting package to NCG),

Italian Association for Cancer Research (AIRC) Investigator Grant (IG) 27101 to NCG Italian Association for Cancer Research (AIRC) Three-year fellowship“MilanoMarathon - oggicorroperAIRC” - Rif. 22461 to MC, MC was a PhD student within the European School of Molecular Medicine (SEMM).

## Author contributions

MC, NCG and PM conceptualized the research project.

MC and PM developed the computer codes and algorithms.

MC, PM, and NCG. analyzed the data.

MC, NK, TD, CM, MG, and PM performed the experiments.

FI analyzed the RNAseq data

NCG and GP provided the resources. MC wrote the original draft.

MC, PM, and NCG reviewed and edited the manuscript.

NCG supervised the research activity.

## Data and materials availability

All data are available in the main text or the supplementary materials.

