## Supplementary material for "Unraveling Glioblastoma Heterogeneity: Introducing SP2G Method for Identifying Invasive Sub-Populations"

Michele Crestani *et al.*

**This PDF file includes:**

Figs. S1 to S6
Movies S1 to S6
Data S1 to S3

**Other Supplementary Materials for this manuscript include the following:**

Movies S1 to S6
Data S1 to S3
Supplementary-Appendix 1

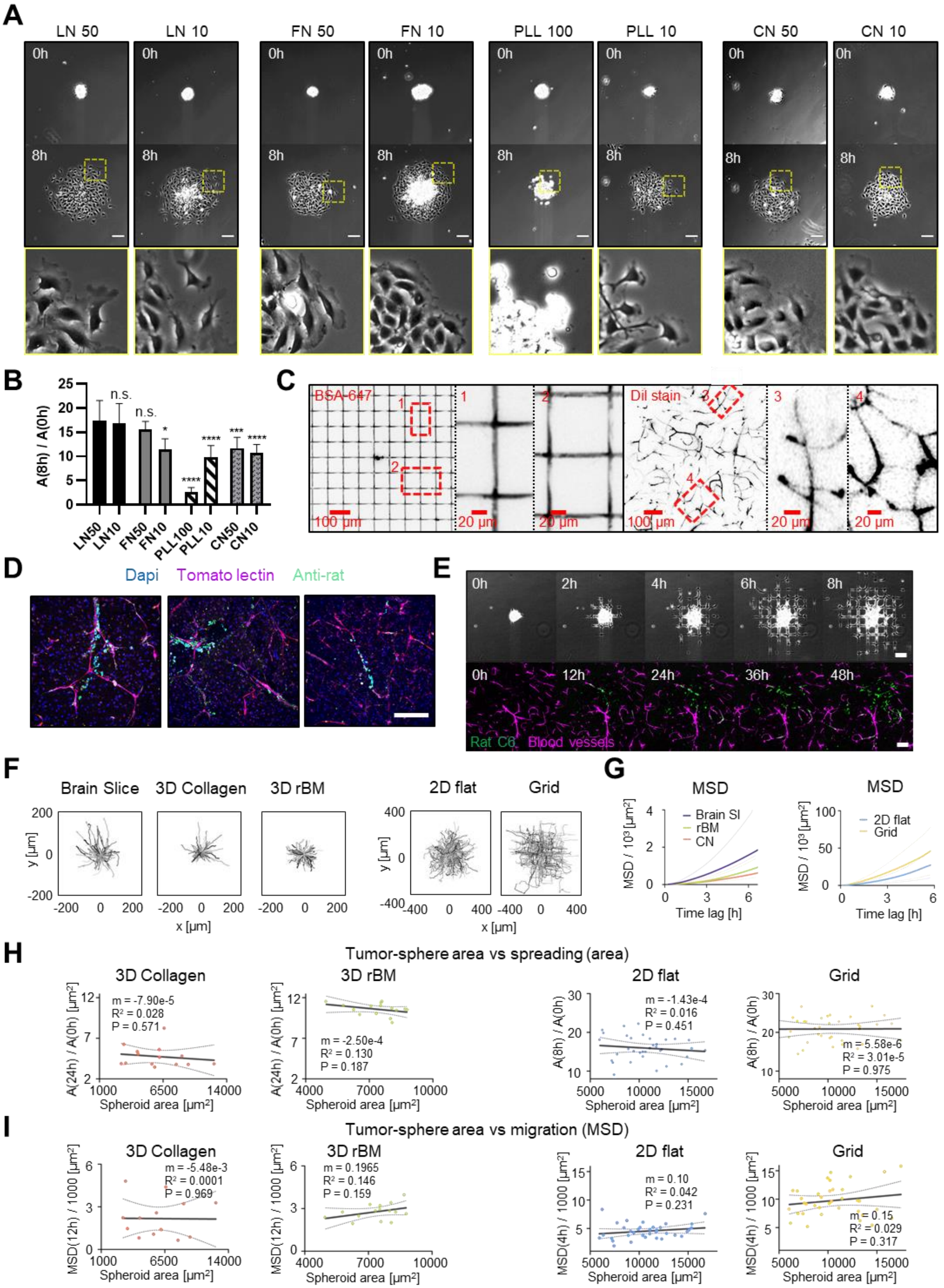
A
LN 50
LN 10
FN 50
FN 10
PLL 100
PLL 10
CN 50
CN 10
0h
8h
B
A(8h) / A(0h)
n.s.
LN50
LN10
FN50
FN10
PLL100
PLL10
CN50
CN10
C
BSA-647
DiI stain
100 µm
20 µm
D
Dapi
Tomato lectin
Anti-rat
E
Rat C6
Blood vessels
F
Brain Slice
3D Collagen
3D rBM
2D flat
Grid
y [µm]
x [µm]
G
MSD
Brain Sl
rBM
CN
2D flat
Grid
Time lag [h]
H
Tumor-sphere area vs spreading (area)
Spheroid area [µm²]
m = -7.90e-5
R² = 0.028
P = 0.571
m = -2.50e-4
R² = 0.130
P = 0.187
m = -1.43e-4
R² = 0.016
P = 0.451
m = 5.58e-6
R² = 3.01e-5
P = 0.975
I
Tumor-sphere area vs migration (MSD)
m = -5.48e-3
R² = 0.0001
P = 0.969
m = 0.1965
R² = 0.146
P = 0.159
m = 0.10
R² = 0.042
P = 0.231
m = 0.15
R² = 0.029
P = 0.317

**Fig. S1. SP2G mimics glioblastoma invasion on brain blood vessels.**

(**A-B**) Rat C6 glioma cells cultured as spheroids were seeded on glass bottom dishes coated with laminin (LN), fibronectin (FN), Poly-L-Lysine (PLL) and collagen (CN) at 10, 50 or 100 µg/ml and imaged for 8h. (**A**) First and last images of the movies (upper panels) and zoom at 8h (lower panels). (**B**) Quantification of spheroid spreading (area at 8 h over the initial area). Error bars are SD. n = 10, 7, 8, 6, 8, 8, 8, 8, spheroids. Each condition was compared to the LN50 condition. (**C**) Comparison of gridded micropatterns (stained with fluorescent BSA, left) with the mouse brain vasculature (stained with Dil dye, right). (**D**) Mouse brain slices invaded by rat C6 glioma cells for 48h, fixed and stained with DAPI, tomato lectin (blood vessels) and anti-rat antibody. (**E**) Montage extracted from time-lapse movies of C6 spheroid spreading on gridded micropatterns (top) and in brain slice (bottom). (**F**) Cell trajectories of C6 single cells migrating in brain slice, collagen gel, reconstituted basement membrane (rBM), on laminin-coated dishes, and gridded micropatterns. n = 80, 95, 90, 215, 215 tracks; 5 to 7 tracks per spheroid, each dot is a cell. (**G**) Mean squared displacement (MSD) plots obtained from single cell tracks in brain slice, collagen gel, rBM, 2D flat and grid (n = 80, 95, 90, 216, 216 tracks; 5 to 7 tracks per spheroid. n = 2, 2, 2, 6, 6 independent experiments ). (**H-I**) Spheroid spreading (**H**) and MSD (**I**) measured in function of the spheroid area at the initial time. Line slope (m), coefficient of determination (R2) and p-value (P) of the linear regression are reported. n = 14, 15, 35, 35 spheroids. Dashed lines are S.D.

**A.** Migration area, A(t). *Semi-automated step.*

Place raw data in a folder. Raw data can be:
- Several bio-format files (e.g. 1 per condition)
- .tif files (1 per spheroid) placed in a sub-folder

Run code "*Main: Polygon Tracking and Segmentation*". Inputs: raw data. Outputs are:

1 folder per raw data file, in which 1 subfolder per sample contains:
- Binarized grid
- Binarized cells
- Polygonal shape
- Binarized grid nodes
- .avi movie
- .txt file (debugging)

- Run code "*Average Polygon*". Input: *polygonal shape.*
- Output: graphical representation for A(t)

**B.** Diffusivity, D = δA(t) / δt; Boundary speed, $v(t) = D(t) / (2\sqrt{(A(t))})$. *Automated step.*

Run code "*Polygon Area Measurement*". Input: *polygonal shape.* Outputs, in the same .csv file :
- Migration areas A(t) (absolute and relative to $t_0$)
- Smoothed A(t)
- Diffusivity D(t)
- Boundary speed v(t)

**C.** Extrapolate τ and Δ from mean A(t) and $v(t)$. *Semi-automated/User step.*

- τ normalizes over migration area. It is the time step when the mean A(t) reaches the lowest A(final_time) among the experimental conditions to compare.
- Δ normalizes over boundary speed. It is the time window taken by the cells to travel 100 µm. If Δ < 8h, cells are considered motile. Δ must be lower than τ.

*See supplementary file 1, it can be utilized as template.*

**D.** Collective migration and Directional Persistence. *Automated step.*

Run code "*Collective migration and Directional Persistence*". Input: *binarized cells*. Output: Collective migration and Directional Persistence values in .csv file (mean and indexed by spheroid name. All values indexed by time point.)

**E.** Hurdling. *Semi-automated step.*

Run code "*Hurdling*". Input: *binarized cells* and *binarized grid*. Outputs:
- Average Intensities (1 value per square) + Mean Average Intensity (1 value) in .csv file, for relative hurdling calculation.
- Average intensities (1 value per square) normalized over the maximum value of the dataset in the same .csv file, for visualizing the cumulative distribution.

All values indexed by time point. *See supplementary file 2.*

**F**

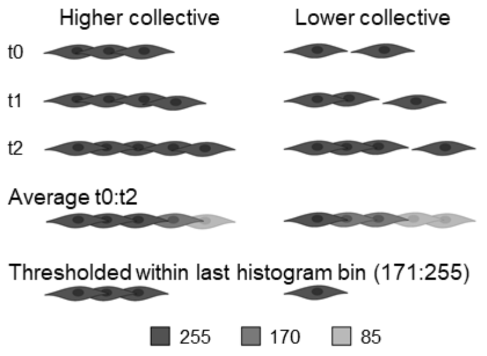


**G**

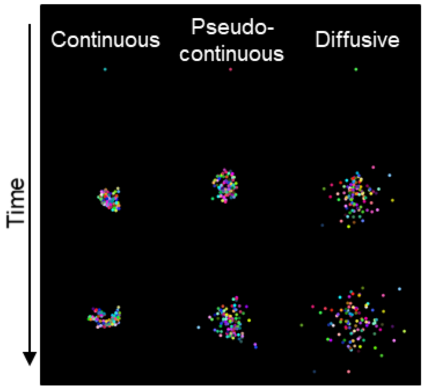


**H**

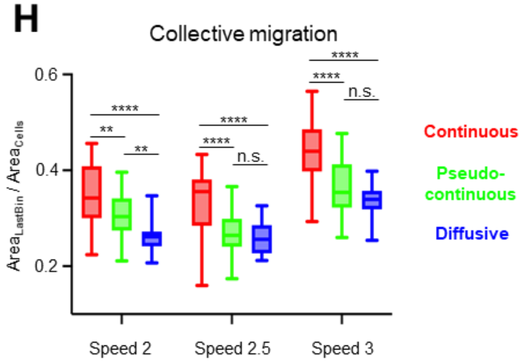


**I**

| Speed | Diffusion | τ−Δ | Δ |
|---|---|---|---|
| 2 | Continuous | 285 | 15 |
| | Pseudo-cont | 105 | |
| | Diffusive | 49 | |
| 2.5 | Continuous | 93 | 12 |
| | Pseudo-cont | 69 | |
| | Diffusive | 31 | |
| 3 | Continuous | 69 | 10 |
| | Pseudo-cont | 49 | |
| | Diffusive | 20 | |

**Fig. S2. SP2G: from images to quantitative data.** (**A-E**)Table summarizing SP2G quantitative outputs. (**F**) Schematic representation of the rationale allowing collective migration quantification. Average images obtained from highly collective cells (left) will return higher values and form higher area when thresholded, compared to low collective cells (right). (**G-I**) SP2G outputs obtained with simulated data using 100 round particles (radius=5 pixels) diffusing with full constraint (Continuous, 100% probability of being attached to a neighbor), partial constraint (Pseudo-continuous, 90% probability of being attached to a neighbor), or no constraint (Diffusive, simple diffusion), over 300 time frames at 3 speed regimes, with mean speed of 2, 2.5, 3 pixels per frame and standard deviation of 2, 2.5, 3 pixels, respectively (9 conditions total). (**G**) Example of output obtained with speed = $3 \pm 3$ pixels per frame. (**H**) Collective migration output obtained in the 9 conditions. (**I**) Table highlighting each simulated condition, their corresponding $\Delta$ ($\Delta$ was chosen so as its product with speed equals 30) and $\tau - \Delta$. n=30 simulations per condition.

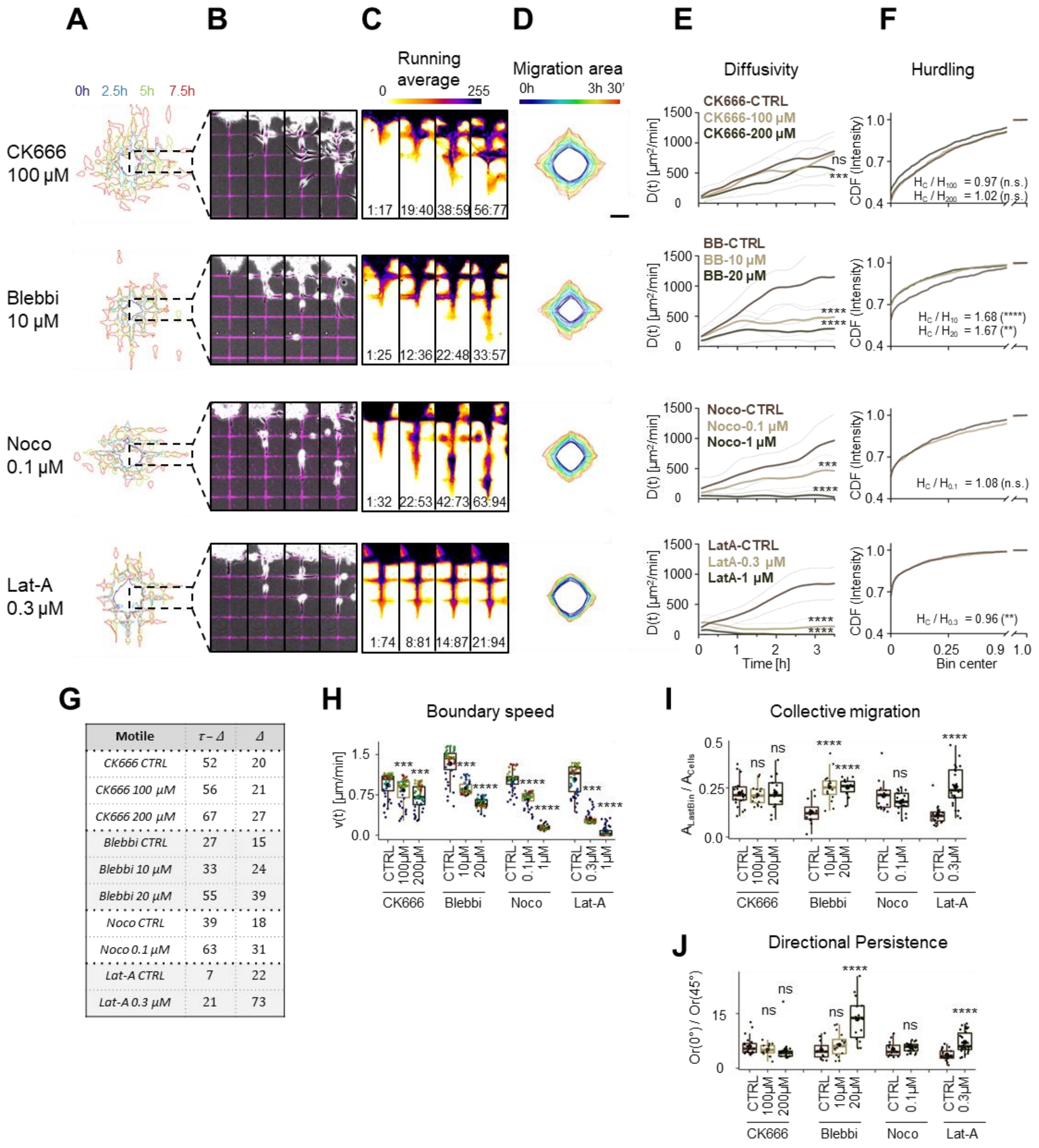


| Motile | τ − Δ | Δ |
|---|---|---|
| CK666 CTRL | 52 | 20 |
| CK666 100 µM | 56 | 21 |
| CK666 200 µM | 67 | 27 |
| Blebbi CTRL | 27 | 15 |
| Blebbi 10 µM | 33 | 24 |
| Blebbi 20 µM | 55 | 39 |
| Noco CTRL | 39 | 18 |
| Noco 0.1 µM | 63 | 31 |
| Lat-A CTRL | 7 | 22 |
| Lat-A 0.3 µM | 21 | 73 |

**Fig. S3. SP2G quantifies cell migratory tactic changes upon cytoskeleton drugs.** Patient-derived-spheroids, NNI-21, were seeded on fluorescent gridded micropatterns, imaged for 8h in presence of each drug, and analyzed as indicated in fig. 2 (n= 16, 14, 16, 14, 20, 12, 26, 16 spheroids for CK666 100 and 200 µM, blebbistatin 10 and 20 µM, Nocodazole 0.1 and 1 µM, latrunculin-A 0.3 and 1 µM respectively and n = 25, 12, 16, 21 for their respective control, n = 2 independent experiments per condition). (**A-C**) Cellular edges at 4 time points (**A**), corresponding overlays of the phase contrast and the fluorescent grid images (**B**) and corresponding running average (RA) (**C**). Δ is indicated at the bottom of each panel. (**D**) Averaged polygons visualizing migration area. (**E**) Diffusivity over 3.5 h. Dashed lines are standard deviation. (**F**) Hurdling visualized as the Cumulative Distribution Function (CDF) of the normalized mean intensity of the grid squares. (**G**) $\Delta$ and $\tau$ - $\Delta$ in motile cells. (**H**) Mean boundary speed over 3.5 h. Each dot represents a time-point and is color-coded as in D. (**I-J**) Collective migration (**I**) and directional persistence (**J**) for the motile cells. Each dot represents a spheroid. Each treated condition was compared against the control. Time and image intensity are color-coded as indicated.

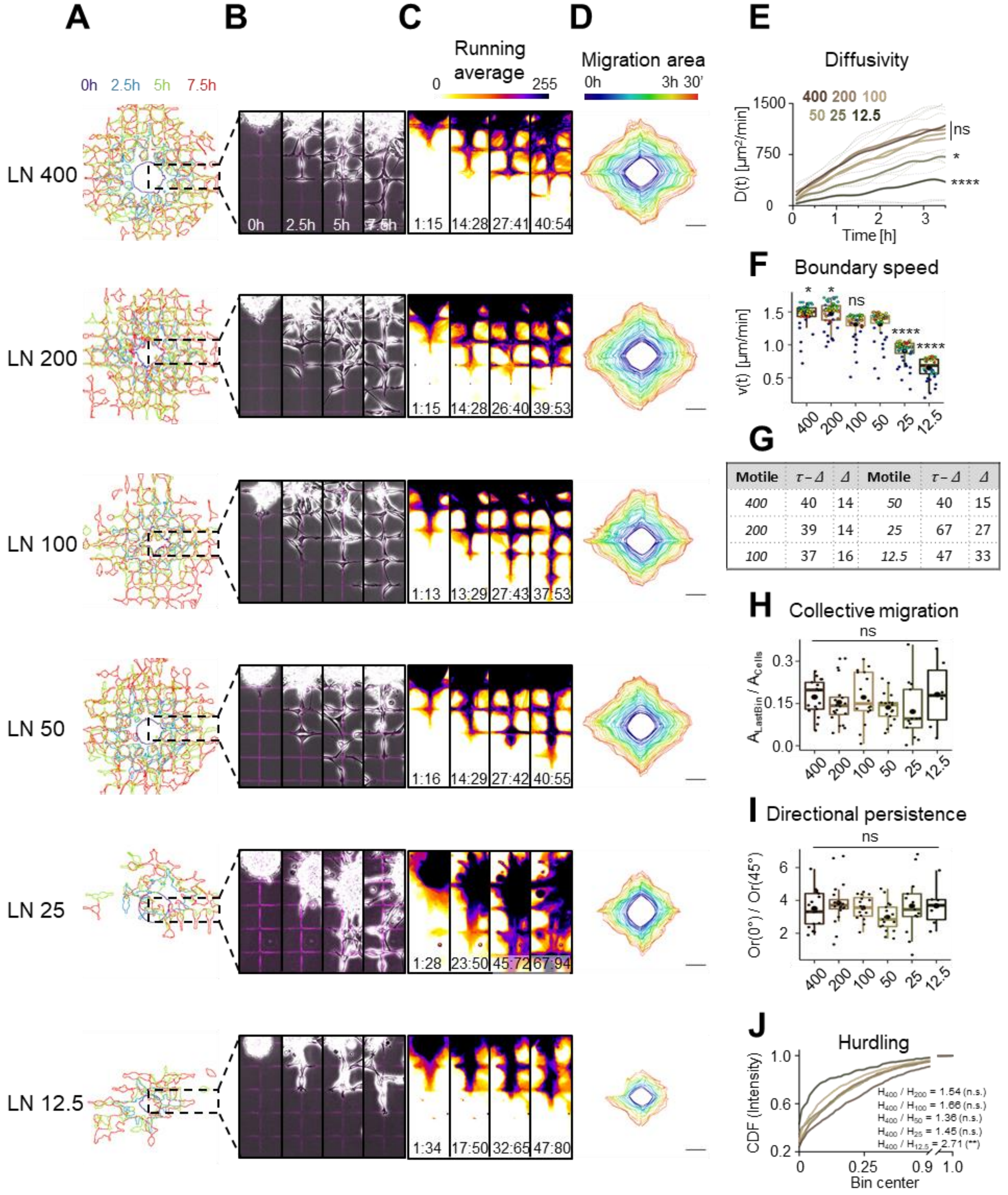


**Fig. S4. SP2G detects cell migratory tactic changes upon variations of laminin concentration.** Analysis of NNI-21 spheroid spreading on gridded micropatterns printed with laminin (LN) at 400, 200, 100, 50, 25, 12.5 μg/ml (n = 17, 20, 16, 17, 13, 6 spheroids, n = 3 independent experiments per condition). (**A-C**) Cellular edges (**A**) and corresponding overlays of the phase contrast and the fluorescent grid images (**B**) at 4 time points and corresponding running average (RA) (**C**). (**D**) Averaged polygons visualizing migration area. (**E**) Diffusivity over 3.5 h. (**F**) Mean boundary speed over 3.5 h. Each dot represents a time-point and is color-coded as in D. (**G**) $\Delta$ and $\tau$ - $\Delta$ in motile cells. (**H-I**) Collective migration (**H**) and directional persistence (**I**) for the motile cells. Each dot represents a spheroid. (**J**) Hurdling visualized as the Cumulative Distribution Function (CDF) of the normalized mean intensity of the grid squares. Time and image intensity are color-coded as indicated. Each condition was compared against LN 50 μg/ml.

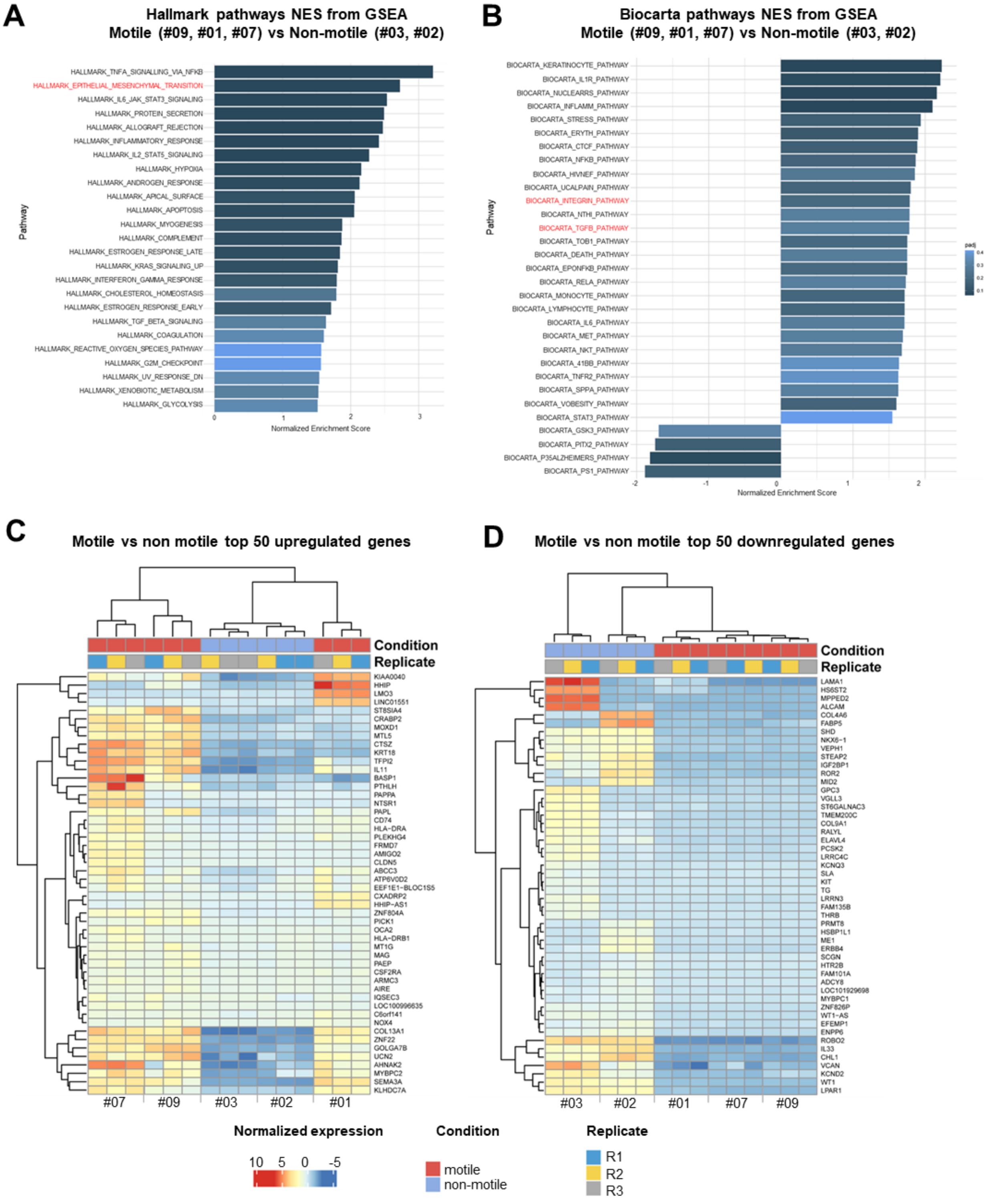


**Fig. S5. Intra-patient heterogeneity in migration ability is correlated with specific molecular signatures.** (**A-B**) GSEA of differentially expressed genes in the motile group (Clones #01, 07, 09) compared to the non-motile group (Clones #02, 03), performed using Hallmark (**A**) and Biocarta (**B**) gene sets from the GSEA molecular signature database (https://www.gsea-msigdb.org/gsea/msigdb ). (**C-D**) Hierarchically clustered heatmaps of the 50 most upregulated genes (**C**) and the 50 most down-regulated genes (**D**) in the motile group compared to the non-motile group.

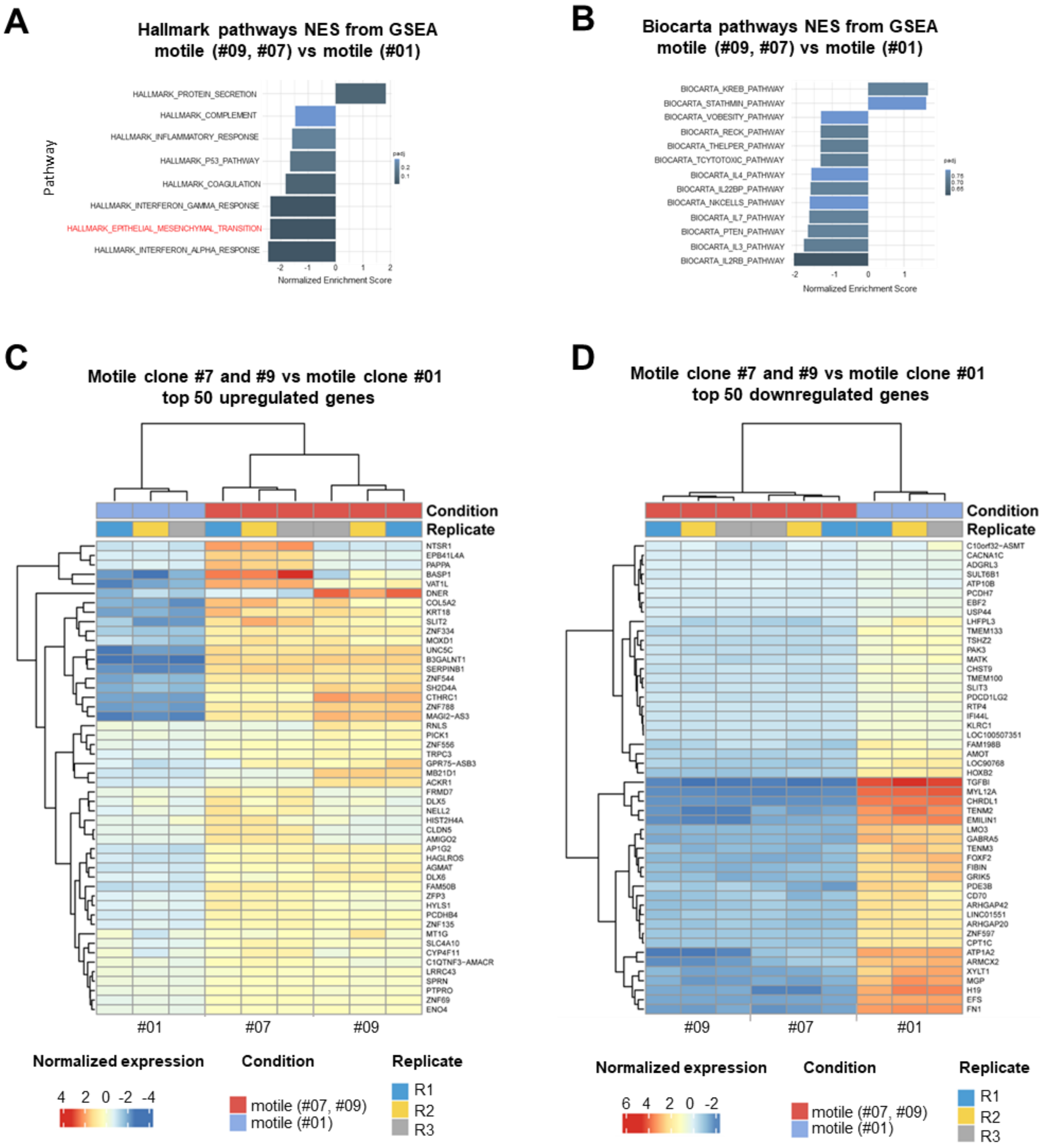


**Fig. S6. Intra-patient heterogeneity in migration modes is correlated with specific molecular signatures.** (**A-B**) GSEA of differentially expressed genes between the 2 motile groups performed using Hallmark (**A**) and Biocarta (**B**) gene sets from the GSEA molecular signature database (https://www.gsea-msigdb.org/gsea/msigdb). The motile clones # 07, 09 were compared to the motile clone #01. (**C-D**) Hierarchically clustered heatmaps of the 50 most upregulated genes (**C**) and the 50 most down-regulated genes (**D**) in the motile clones #07, 09 compared to the motile clone #01.

**Movie S1.**

Concatenated timelapse movies of spreading C6 spheroids. 1) brain slices; 2) 3D collagen and matrigel; 3) 2D flat and gridded micropatterns; 4) representative single cells moving in each setup.

**Movie S2.**

Tutorial for SP2G macro suite

**Movie S3.**

SP2G known migration phenotypes. 1) SP2G of NNI-11, NNI-21, NNI-24; 2) SP2G tracking; 3) SP2G averaging; 3) SP2G projected averaging; 4) running average for the motile NNI-21 and NNI-24.

**Movie S4.**

SP2G drug treatments (NNI-21). Top: SP2G tracking, bottom: SP2G averaging. 1) CK666; 2) Blebbistatin; 3) Nocodazole; 4) Latrunculin-A.

**Movie S5.**

SP2G in the GBM7 patient derived cell line. 1) Heterogeneous bulk; 2) intra-patient heterogeneity of the 5 clones; 3) SP2G tracking; 4) SP2G averaging; 5) SP2G projected averaging; 6) running average for the motile #09, #01, #07; 7) spreading spheroids in brain slices of the 5 clones.

**Data S1. (separate file)**

Calculations of Tau, Delta, and mean boundary speed starting from the time trend of the relative area occupied by cells. Imposed parameters: motility threshold (does the invasive front travel faster than a given space - in our case 100 microns - in 8 hours? yes/no), frame rate, and total number of frames.

**Data S2. (separate file)**

Calculations of Average Hurdling and Relative hurdling starting from the raw distribution of intensities in the passivated areas of the grid.

**Data S3. (separate file)**

Details of all the statistical analyses.

**Supplementary-Appendix 1. (separate file)**

Step by step manual for running SP2G macro suite.
